# A microexon confers functional divergence of the actin nucleator Arp2/3

**DOI:** 10.1101/2025.10.12.681909

**Authors:** Jordan Powell, Manuela Sophia Palafox, Courtney M. Schroeder

**Affiliations:** Department of Pharmacology, UT Southwestern Medical Center, Dallas, TX

**Author notes:** Co-first authors (Jordan Powell, Manuela Sophia Palafox).

**Keywords:** cytoskeleton, actin, Arp2/3, evolution, microexon, splice variants, sperm development

## Abstract

*Arp2*, a subunit within the actin nucleator Arp2/3, encodes two splice variants that differ by five residues in the D-loop, a critical element for catalyzing actin polymerization. It is unknown if this alternative exon, or “microexon,” impacts Arp2/3 function. Here, we find that the microexon has been evolutionarily retained for over 600 million years yet varies in sequence. We investigate the unique microexon in *Drosophila* Arp2 and show that the longer “Arp2L” variant is expressed, though less than the short variant, “Arp2s.” We purify recombinant *Drosophila* Arp2/3 containing Arp2s or Arp2L, and although we do not detect differences in actin polymerization bulk assays, the microexon attenuates inhibition by Arp2/3 inhibitors *in vitro*, suggesting intrinsic differences. We generate flies expressing only *Arp2s* or *Arp2L*, and while both splice variants rescue *Arp2*-knockout lethality, they functionally diverge in late sperm development due to the microexon sequence, not the length. Overall, our findings demonstrate that the Arp2 microexon alters Arp2/3 function in certain contexts.

## Introduction

Nucleators spatially and temporally control actin polymerization *in vivo*. Arp2/3 is a highly conserved actin nucleator that arose early in eukaryotic evolution (Goodson and Hawse, 2002; Muller et al., 2005) and uniquely produces branched actin networks (Amann and Pollard, 2001; Mullins et al., 1998). Branched networks provide pushing forces in numerous processes, including cell motility (Pollard and Borisy, 2003; Suraneni et al., 2012; Wu et al., 2012) and vesicle trafficking (Chakrabarti et al., 2021). The name of the Arp2/3 complex highlights the two subunits that are close relatives of actin: actin-related proteins (Arps) 2 and 3, which form the actin branch’s base (Chou et al., 2022; Ding et al., 2022). Arp2, Arp3, and the remaining five scaffolding subunits, Arpc1-5, dock onto a pre-existing filament and generate a branch at ∼70° from the “mother” filament (Mullins et al., 1998) upon activation by a nucleation-promoting factor (NPF) (Gautreau et al., 2022; Machesky et al., 1999).

Several of the Arp2/3 subunits have diversified in the form of paralogs in specific species. For example, humans encode two paralogs of Arp3, Arpc1, and Arpc5 (Abella et al., 2016). These isoforms diverge in rates of actin polymerization or network disassembly (Abella et al., 2016; Galloni et al., 2021) and are non-redundant *in vivo* (Kahr et al., 2017; Kuijpers et al., 2017; Nunes-Santos et al., 2023; Randzavola et al., 2019; Sindram et al., 2023; Vasconcellos et al., 2024). Flies uniquely encode paralogs of the subunits Arp2 and Arpc3 that play tissue-specific roles (Hudson and Cooley, 2002; Schroeder et al., 2020; Stromberg et al., 2023). Therefore, evolution has modified the components of the complex to adapt for multiple contexts.

The Arp2 D-loop is a prime site of sequence divergence among paralogs and splice variants. The D-loop is a disordered region in subdomain 2 of the actin fold found in all Arps (Dominguez and Holmes, 2011). It contacts the subunit Arpc3 (Chou et al., 2022; Ding et al., 2022) (Fig. 1A), and this interaction is critical for complex activation (Ding et al., 2022). Despite its importance, the D-loop has diverged in sequence in clade-specific Arp2 paralogs and has specialized Arp2/3 function for actin branching in the testis (Stromberg et al., 2023). Furthermore, Arp2 shows variation in this region in one of two splice variants (Ozturk-Colak et al., 2024; Perez et al., 2025). The longer Arp2 splice variant includes an alternative exon that encodes only 5 amino acids, a “microexon.” Microexons typically encode 1-10 amino acids but can reach up to 17 residues (Li et al., 2015; Ustianenko et al., 2017) and have been identified in humans, *C. elegans,* and *D. melanogaster* (Choudhary et al., 2021; Irimia et al., 2014; Pang et al., 2021; Torres-Mendez et al., 2022). Some microexons alter unstructured loops or sequences within protein-interaction domains and are often developmentally regulated and tissue-specific (Irimia et al., 2014; Li et al., 2015). The Arp2 microexon may augment mechanisms of Arp2/3 regulation, yet its functional relevance is unknown.

**Figure 1:**
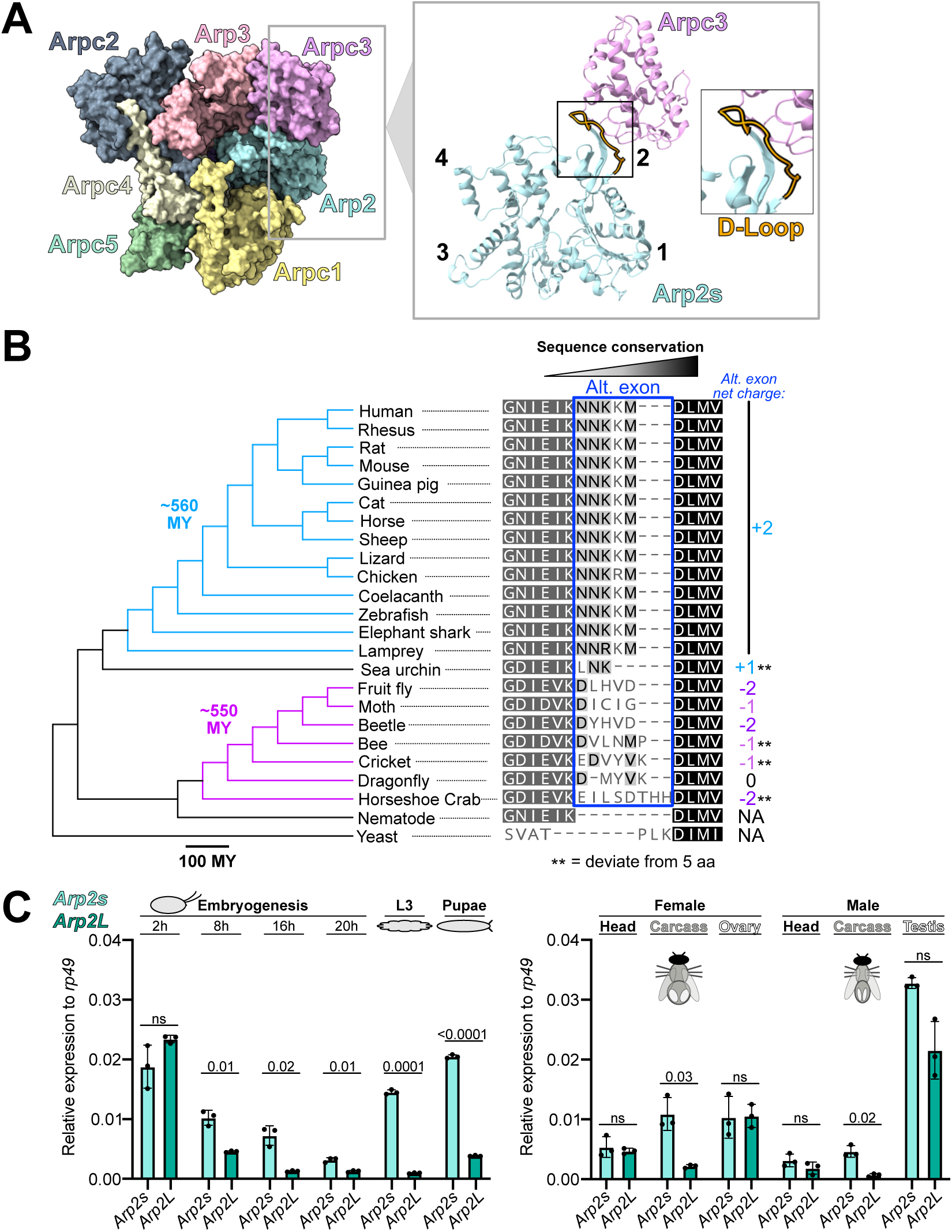
An alternative exon in the Arp2 D-loop diverged in sequence throughout eukaryotic evolution and exhibits a restricted expression profile in *Drosophila*. **A)** Cryo-EM structure of Arp2/3 (PDB 7TPT) (Ding et al., 2022). Structures of Arp2s and Arpc3 are shown alone with an inset highlighting the interaction between the D-loop (orange) in subdomain 2 of Arp2s with Arpc3. **B)** Alignment of a portion of Arp2L sequences from 22 species with two Arp2 sequences from *C. elegans* and *S. pombe*, which do not encode a microexon. Overall charge of the microexons shown on the right. Species tree on the left shows vertebrate lineages in blue and insects in magenta. Shading of residues indicates relative conservation (100% conserved in black, least conserved in white). Microexon (boxed in blue) is strictly conserved in vertebrates but diversified in invertebrates. **C)** Quantitative real time-PCR analysis of *Arp2s* and *Arp2L* expression throughout *Drosophila* development (with four timepoints throughout embryogenesis) and the head, germline, and remaining carcass from adult females and males. Expression is normalized to *Rp49* within each sample (*Rp49* is lower in the testis, resulting in higher *Arp2* relative expression in the testis versus other adult tissues). Statistical test: unpaired t-test with Welch’s correction. Fly diagrams indicate adult tissues analyzed. *Arp2L* expression is restricted in expression.

Here, we show that evolution has retained the Arp2 microexon for over 600 million years, yet its sequence diverges among invertebrates. We test the *Drosophila* Arp2 microexon for any impact on Arp2/3 function. We find that *Drosophila* Arp2/3 with “Arp2s” (short variant) or “Arp2L” (long variant encoding the microexon) exhibits similar polymerization rates in bulk assays with the NPF WASP, yet Arp2L complexes exhibit less inhibition by Arp2/3-targeting drugs, indicating the microexon can alter Arp2/3 function *in vitro*. We then test for functional differences *in vivo*. While adult flies expressing only Arp2s or Arp2L are viable, flies expressing only Arp2L exhibit defects in actin structures in late sperm development. We discover that the microexon sequence underlies this phenotype, rather than the five-residue increase in the D-loop. Despite these fitness costs, our evolutionary experiments suggest that encoding Arp2L provides an overall fitness advantage. These findings demonstrate that the microexon impacts Arp2/3 function *in vitro* and *in vivo* and is a potential source of Arp2/3 regulation.

## Results and Discussion

### An *Arp2* microexon diversifies in sequence among invertebrates and is expressed

To gain insight into how widespread the Arp2 microexon is across species and to assess its sequence conservation, we compared the *Arp2* sequence from yeast to human (Fig. 1B, Fig. S1A). We found that all surveyed vertebrates have retained the microexon (∼560 million years of evolution), including one of the earliest groups of vertebrates—lampreys (Xu et al., 2016). The vertebrate sequence is also stringently conserved, except for the rare substitution of a Lys to Arg.

Interestingly, the microexon dramatically varies in sequence among invertebrates. While vertebrate sequences maintain an overall positive charge, invertebrate sequences are negative or neutral. Some species’ microexon diverges in length. For example, sea urchin encodes only three residues (Fig. 1B), and bees encode three splice variants, with one reaching 12 residues long (Table S1). Because the invertebrate sequences are unusual, we verified their expression using publicly available RNA expression databases (Table S1). We conclude that the microexon is stringently conserved in vertebrates, yet it diversified in sequence and length among invertebrates, suggesting species specialization. Yeast and nematodes do not encode the microexon (Fig. 1B). Thus, its evolutionary history suggests that the microexon was present in the last common ancestor of insects and vertebrates approximately 686 million years ago (Fig. 1B) (Kumar et al., 2022), yet its origin may be older.

We focused on the long *Arp2* splice variant (“*Arp2L*”) in *D. melanogaster* to probe the biological importance of the microexon. Despite sharing the same length as the vertebrate microexon, the *Drosophila* microexon drastically differs in sequence and is negatively charged (Fig. 1B, Fig. S1A), which is conserved among all Drosophilids (Fig. S1B). First, we investigated whether *Drosophila Arp2L* is even expressed, comparing its expression with that of the splice variant lacking the microexon (“*Arp2s*”) across different developmental stages and adult tissues. We extracted mRNA from wildtype embryos (at multiple developmental time points), larvae, and pupae. We found that *Arp2s* is expressed throughout development, and although *Arp2L* is highly expressed in early embryogenesis, its expression decreases in late embryogenesis and the larval and pupal stages (Fig. 1C, Table S2). *Arp2L* is expressed comparably to *Arp2s* in the ovary and head. In contrast, female and male somatic tissue (excluding the head and germline) exhibits significantly less *Arp2L* expression than *Arp2s*. Males also show less *Arp2L* than *Arp2s* mRNA in the testis, though not significant due to variability (Fig. 1C, Table S2). In sum, *Arp2s* is ubiquitously expressed, yet *Arp2L* is restricted in expression.

### Arp2s and Arp2L exhibit different degrees of inhibition by Arp2/3-targeting drugs *in vitro*

Because the D-loop is critical for actin polymerization (Ding et al., 2022), we hypothesized that the microexon in Arp2L directly impacts the rate of polymerization. All Arp2/3 structures to date include Arp2s, giving little insight into how the microexon may structurally impact Arp2/3 activity (Ding et al., 2022; Robinson et al., 2001; Zimmet et al., 2020). An AlphaFold 3-predicted structure (Abramson et al., 2024) of *Drosophila* Arp2L in the complex shows maintenance of the D-loop-Arpc3 interaction (Fig. S2C) yet exhibits low confidence in modeling the microexon.

To directly compare Arp2s and Arp2L activity *in vitro*, we affinity-purified recombinant *Drosophila* Arp2/3 via a FLAG-tag encoded on the C-terminus of Arpc3, similar to a human Arp2/3 study (Zimmet et al., 2020). *Drosophila* uniquely encodes two isoforms of the Arpc3 subunit: Arpc3A (orthologous to human Arpc3) and Arpc3B, found only in *Drosophila* (Hudson and Cooley, 2002) (Fig. 2A). We purified all four combinations of subunits in the Arp2/3 complex and verified the subunit composition of all purified complexes by intact mass spectrometry (Fig. 2B, Table S2). Because Arp2/3 requires an NPF to catalyze actin polymerization (Higgs et al., 1999; Machesky et al., 1999), we purified the activating domain of the *Drosophila* NPF WASP. The activating domain of the WASP protein family is the conserved “VCA” domain, composed of a verprolin-homology domain, the cofilin- (or central-) homology domain, and the acidic domain (Takenawa and Suetsugu, 2007). We purified a dimeric GST-tagged WASP VCA domain, as a dimer is a more potent activator (Padrick et al., 2008).

**Figure 2:**
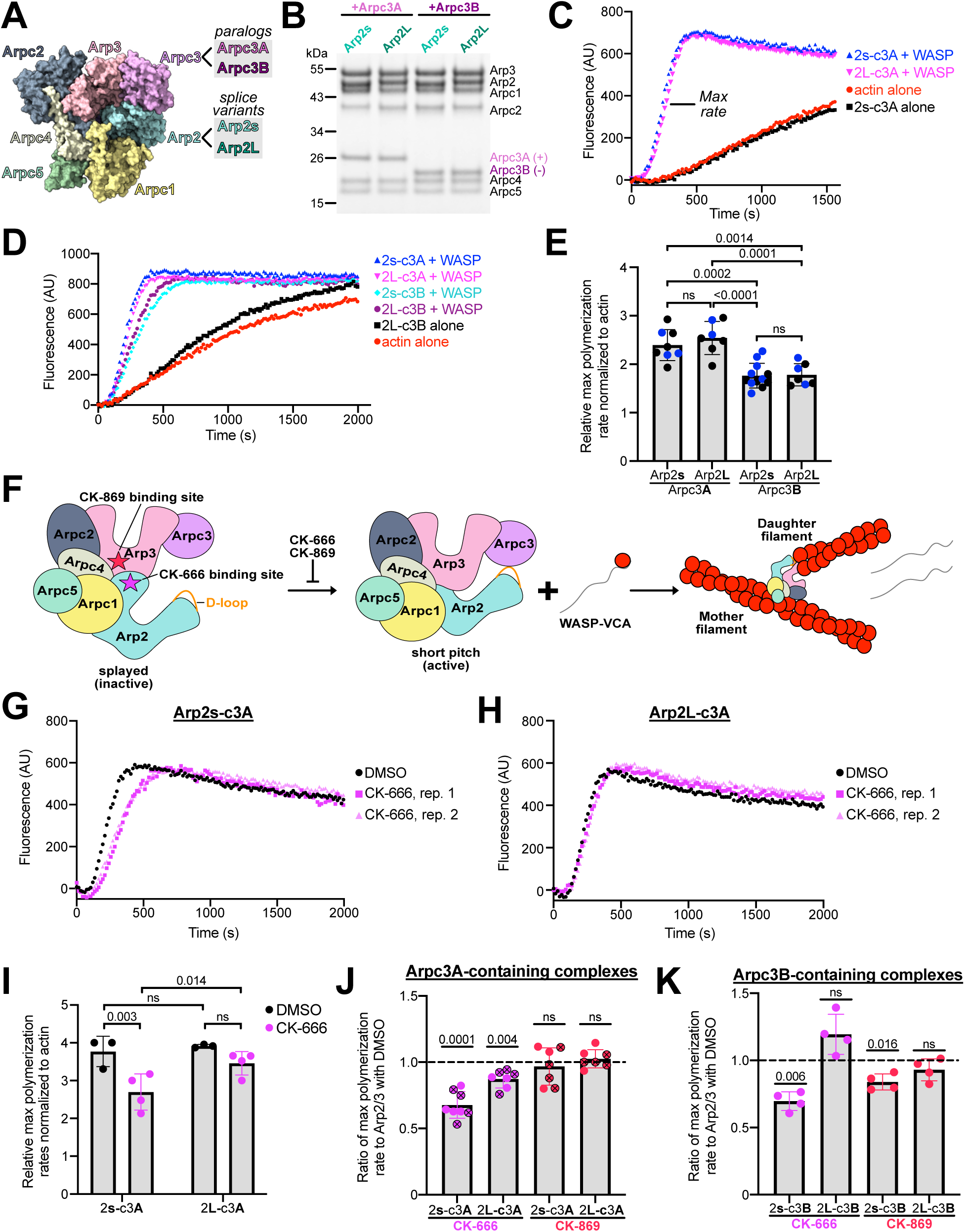
*Drosophila* Arp2s and Arp2L polymerize actin at similar rates *in vitro*, but the microexon attenuates inhibition by Arp2/3-targeting drugs. **A)** Arp2/3 structure (PDB 7TPT) (Ding et al., 2022). Paralogs of Arpc3 and splice variants of Arp2 are noted. **B)** Protein gel showing purified recombinant *Drosophila* Arp2/3 proteins diluted at an equal concentration for polymerization assays in C-D. Bands are labeled, and overall charges of FLAG-tagged Arpc3A and Arpc3B are indicated. Arpc3B, although the same molecular weight as Arpc3A, migrates farther due to its negative charge. **C-D)** Representative graphs showing a time course of actin polymerization, with 2 µM actin (10% pyrene labeled), 20 nM Arp2/3, and 100 nM WASP (GST-VCA domain). Increasing fluorescence indicates increasing actin polymerization, and the maximum polymerization rate is at the curve’s inflection point. **E)** Superplot showing the ratio of the maximum slope of a polymerization curve (i.e., maximum rate, as noted in panel C) divided by the maximum slope of the curve for actin alone. Color of datapoints correspond to a distinct batch of purified actin used in the assay. Statistical test: a one-way ANOVA with a Tukey’s multiple comparisons post-hoc test. **F)** Schematic depicting the binding sites of CK-666 and CK-869 in the Arp2/3 complex and how they inhibit the conformational change from the inactive splayed form to the short-pitch active form of Arp2/3, preventing WASP-stimulated actin branching. **G-H)** Representative graphs showing actin polymerization by WASP-stimulated Arp2s-Arpc3A (G) or Arp2L-Arpc3A (H) over time as in C-D. Reactions included either DMSO (negative control) or 100 µM CK-666. Two replicates shown. **I)** Plot showing the ratio of the maximum slope of a polymerization curve divided by the maximum slope of the curve for actin alone as in panel E. Statistical test: a two-way ANOVA with a Tukey’s multiple comparisons post-hoc test comparing the means of each complex with and without CK-666. **J-K)** Plots showing the ratio of the maximum slope of a polymerization curve (Arp2/3 plus CK-666 or CK-869) divided by the maximum slope of the curve for Arp2/3 plus DMSO. Statistical test: a one-sample ratio t-test with the hypothetical value being 1. Panel J is a superplot; datapoints with an “X” correspond to experiments done with a distinct batch of purified actin.

We compared the actin polymerization rates of the purified Arp2/3 complexes using pyrene-labeled skeletal muscle actin, which increases in fluorescence upon polymerization. We detected rapid actin polymerization by Arp2/3 only in the presence of the VCA domain, as expected (Fig. 2C), and we found that Arp2s and Arp2L in complex with Arpc3A showed no difference in WASP-stimulated polymerization rates (Fig. 2C, E). We next tested whether they differed when in complex with Arpc3B and again detected similar polymerization rates (Fig. 2D-E). Despite no differences found between Arp2s and Arp2L, both splice variants displayed slower polymerization with Arpc3B across independent protein purifications (Fig. 2D-E, Fig. S2A-B).

More subtle differences between the Arp2 splice variants may exist that are not apparent in our bulk assay. One difference that arises among human Arp2/3 iso-complexes is drug susceptibility, even when differences in complex composition are distal to drug-binding interfaces (Cao et al., 2024). Thus, we challenged our *Drosophila* Arp2/3 complexes *in vitro* with two Arp2/3-targeting drugs, CK-666 and CK-869, to assess differences in responses between complexes containing Arp2s versus Arp2L when stimulated by WASP. Both drugs prevent the conformational change required for Arp2/3 activation (Hetrick et al., 2013) (Fig. 2F). CK-666 binds a pocket at the Arp2–Arp3 interface distal to the D-loop (Baggett et al., 2012), and CK-869 binds Arp3 (Hetrick et al., 2013) (Fig. 2F, Fig. S2D). Thus, the drug-binding sites do not differ between Arp2s and Arp2L, as they lie outside the microexon. Furthermore, the CK-666 drug-binding site is highly conserved between flies and mammals, while the CK-869 binding interface exhibits some divergence (Fig. S2E). Given where the drugs bind, any observed differences between Arp2s- and Arp2L-containing complexes would be allosterically conferred by the microexon. Using our pyrene assay, we treated *Drosophila* Arp2/3 complexes with 100 µM CK-666 or CK-869, a concentration traditionally used to inhibit Arp2/3 *in vivo* and *in vitro* (Cao et al., 2024). We discovered that Arp2L in complex with Arpc3A exhibited weak inhibition by CK-666 (Fig. 2H-J), whereas Arp2s was significantly inhibited (Fig. 2G, I-J). Neither was inhibited by the Arp3-binding drug CK-869 at this concentration (Fig. 2J, Fig. 2SF). When in complex with Arpc3B, Arp2s still exhibited notable inhibition by CK-666, yet we detected no inhibition of Arp2L (Fig. 2K, Fig. S2G). This difference extends to CK-869, with only Arp2s inhibited by the drug, albeit weakly (Fig. 2K, Fig. S2H). Divergence of the CK-869 binding site between flies and humans may explain the weak inhibition relative to that with CK-666 (Fig. S2E). By increasing the drug concentration to 200 µM, we observed only weak inhibition of Arp2L-Arpc3B by CK-666, and still no inhibition by CK-869 (Fig. S2I). The differences in inhibition between Arp2s and Arp2L indicate that the presence of the microexon attenuates inhibition by Arp2/3-targeting drugs, demonstrating that the microexon confers intrinsic differences *in vitro*.

### Males expressing only Arp2L exhibit defects in testis-specific actin structures

We next explored whether the two splice variants have any functional differences *in vivo*. We adopted a similar approach to that in a previous study, in which we engineered *D. melanogaster* to express only *Arp2s* (Stromberg et al., 2023). We used our *Arp2*-knockout (KO) flies that encode an *attP* docking site in the KO locus on the X chromosome (Stromberg et al., 2023) and inserted the coding sequence of *Arp2L* to compare to flies expressing only *Arp2s* (Fig. 3A, Fig. S1C). Both transgenes are untagged and encode the same 1 kb upstream and downstream regions of endogenous *Arp2*; therefore, they encode the same UTRs and are subject to the same transcriptional control. We also generated lines with *Arp2s* and *Arp2L* encoded on the second chromosome, outside the KO locus, to test if phenotypes are consistent across genetic backgrounds (Fig. S1C).

**Figure 3:**
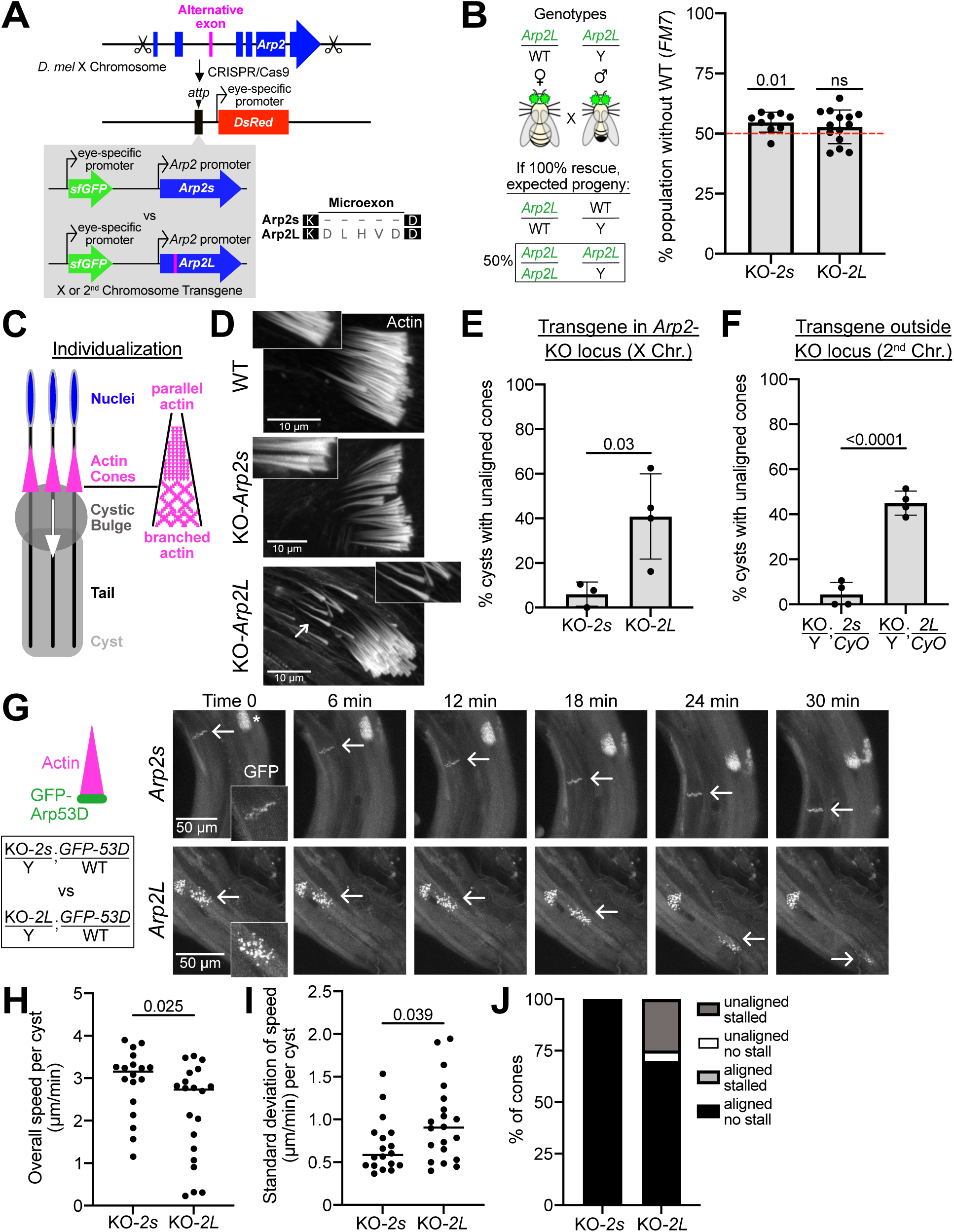
Arp2s and Arp2L are functionally non-equivalent in late sperm development. **A)** CRISPR strategy for generating the *Arp2* knockout (KO) fly line with cut sites in the regions flanking the *D. mel Arp2* coding sequence. *DsRed*, expressed in the eye, replaced *Arp2*. The *attP* site located upstream of *DsRed* was used for site-directed insertion of each *Arp2* splice variant. Transgenes *Arp2s* and *Arp2L* were flanked by 1-kb genomic fragments immediately 5′ and 3′ of the endogenous *Arp2* locus, including the native regulatory elements. A gene encoding *superfolder GFP (sfGFP)* is closely linked to the transgenes to track them by GFP fluorescence in the eye. Transgenes were inserted into the *attP* site of the KO locus on the X chromosome or on the second chromosome of a wildtype fly, then crossed into the KO background. Protein alignment of Arp2s and the Arp2L microexon shown on the right. **B)** Left, crosses to test rescue of *Arp2*-KO zygotic lethality. Heterozygous (Het) females were crossed to hemizygous males. Het females were balanced with *FM7* (denoted as “WT”), which encodes the wildtype *Arp2* locus and confers a distinct eye shape for identification. GFP-positive eyes represent presence of the transgene. Right, percent of total progeny without the WT Arp2 allele (*FM7* balancer) and homozygous for the transgene *Arp2s* or *Arp2L*. Dotted line indicates the expectation based on Mendelian inheritance. Statistical test: one-sample ratio t-test with the hypothetical value being 50%. Number of KO-*Arp2s* progeny was slightly higher than expected, likely indicating a small fitness cost from the *FM7* balancer. Average number of flies counted per replicate: *Arp2s*: 85, *Arp2L*: 73 **C)** Schematic of near-mature sperm undergoing individualization via actin cones. Each cyst includes 64 sperm, and each sperm has one cone. All 64 cones in *D. melanogaster* move synchronously down sperm tails removing excess cytoplasm (“cystic bulge”), encompassing each sperm with its own membrane. Cones contain Arp2/3-generated branched actin networks in the front half and parallel actin filaments in the rear. **D)** Images show a set of actin cones in one cyst of a wildtype testis (*w^1118^* flies), KO-*Arp2s* or KO-*Arp2L* testes. The arrow denotes trailing, unaligned cones in *Arp2L*-expressing males. **E-F)** Quantification of the percent of cysts with unaligned cones. Cones classified as “unaligned” if five or more trail behind the remaining cones within a cyst. Each datapoint represents a single cytological experiment quantifying total individualizing cysts across about 10 males per genotype. Panel E compares flies encoding *Arp2s* or *Arp2L* in the *Arp2*-KO locus. Panel F compares flies expressing only *Arp2s* or *Arp2L* in a different genetic background (*Arp2*-KO flies encoding the Arp2 transgene on the second chromosome balanced with *CyO*). Males encoding only one copy of *Arp2s* or *Arp2L* were analyzed for unaligned cones because WT males encode one copy of *Arp2*. Arp2L-generated cones were frequently unaligned. Statistical test: unpaired t-test with Welch’s correction. **G)** Left, schematic illustrates GFP-Arp53D localized to the leading edge of actin cones, used to visualize actin cone motility in testes *ex vivo*. Males encoding *Arp2s* or *Arp2L* in the *Arp2*-KO locus and GFP-Arp53D on the second chromosome were imaged. A montage of actin cone movement is shown over 30 minutes. Arrows indicate GFP-Arp53D at the leading edge of cones in a single cyst. Asterisk notes the end of a cyst where Arp53D also localizes but was not analyzed. Arp2L-generated cones are unaligned and stalled for over 10 min, eventually moving at a rapid rate. See Videos 1 and 2. **H)** Graph showing the overall speed (µm/min) for a set of cones (64 cones in one cyst). Statistical test: an unpaired t-test with Welch’s correction. **I)** Graph showing the standard deviation of speed (µm/min) per set of cones in a cyst. Standard deviation was taken of the speeds per frame. Statistical test: an unpaired t-test with Welch’s correction. **J)** Graph showing the categorization of motile cones per cyst, including aligned versus unaligned or stalled versus not stalled. Only cones that exhibited motility in at least one frame were analyzed. Number of cysts that were quantified for (6-7 males each): Arp2s: 18 cysts; for Arp2L: 20 cysts. For Arp2s-versus Arp2L-generated cones, stalled cones: p = 0.048; for unaligned cones: p = 0.021. Statistical test: a Fisher’s exact test.

Homozygous *Arp2*-KO flies are embryonic lethal, and in our past study, we found that flies expressing only *Arp2s* can fully rescue *Arp2*-KO lethality with no apparent phenotypes (Stromberg et al., 2023). Here, we investigated whether flies could survive expressing only *Arp2L*. Whether *Arp2L* was encoded in the KO locus or on the second chromosome, we obtained homozygous adults expressing only *Arp2L*, suggesting that Arp2L rescues KO lethality. Using the lines encoding *Arp2L* in the KO locus (“KO-*Arp2L*”), we further quantified the extent of rescue to uncover any fitness cost to zygotic viability. Because *Arp2* is encoded on the X chromosome, males are hemizygous for the *Arp2L* allele. We crossed hemizygous KO-*Arp2L* males to heterozygous KO-*Arp2L* females balanced with *FM7*, a balancer that encodes the WT *Arp2* locus and confers a distinct eye shape for identification (Fig. 3B). If no fitness cost in zygotic viability results from the microexon, ∼50% of the progeny from this cross should include homozygous KO-*Arp2L* females and hemizygous KO-*Arp2L* males, according to Mendelian inheritance (Fig. 3B). Similar to the cross with *Arp2s*-expressing flies, the cross with KO-*Arp2L* flies resulted in approximately 50% of the progeny expressing only Arp2L (Fig. 3B), indicating no cost to zygotic viability.

Having established that flies expressing *Arp2s* or *Arp2L* are viable, we next tested their fertility. KO-*Arp2s* flies were fertile, yet KO-*Arp2L* parents exhibited greatly reduced fertility (Fig. S2J), in which laid embryos reached late embryogenesis but failed to hatch (Fig. S2J, L-N). This developmental phenotype was observed only with the progeny of KO-*Arp2L* females, not males, indicating a maternal effect (Fig. S2K). However, we observed this defect only when *Arp2L* was expressed from the X chromosome, not the second-chromosome transgene (Fig. S2J). Because developmental differences depended on genetic background, we did not further investigate development and instead focused on Arp2s-versus Arp2L-generated actin structures in adult flies to identify differences consistent across genetic backgrounds.

We first examined well-studied Arp2/3-generated structures known as “actin rings” in the ovary. Actin rings allow for the passage of nutrients between the developing oocyte and surrounding “nurse cells.” Upon loss of Arp2/3, actin rings fail to expand in size (Hudson and Cooley, 2002). We measured the size of actin ring diameters across genetic backgrounds and detected no difference in ring size between *Arp2s*- and *Arp2L*-expressing flies, including KO-*Arp2L* females (Fig. S3A-C). We also investigated bristles in the thorax of the adult fly, where *Arp2L* is less expressed than Arp2s (Fig. 1C). Upon loss of Arp2/3, long and short bristles exhibit kinks (Frank et al., 2006; Koch et al., 2012), yet we observed no significant bristle defects between *Arp2s*- and *Arp2L*-expressing flies across genetic backgrounds (Fig. S3D-E).

We next investigated the male germline for differences because, in a previous study, we found that divergence in the D-loop of species-specific Arp2 paralogs led to differences in actin branching in the testis (Stromberg et al., 2023). KO-*Arp2s* flies exhibit no actin defects in the testis (Stromberg et al., 2023); however, here, we investigated whether the same held true for KO-*Arp2L* flies. We investigated a prominent Arp2/3-generated structure: actin cones. Cones separate syncytial developing sperm in *Drosophila* during the final step of sperm maturation (“individualization”). All 64 actin cones within a cyst of individualizing sperm move synchronously down the sperm tail, removing excess cytoplasm and encasing each sperm in its own membrane (Fabrizio et al., 1998; Noguchi and Miller, 2003). Arp2/3-generated branched actin composes the front half of cones and gives cones their fan-like structure (Fig. 3C). Loss of Arp2/3 activity results in needle-like cones that infrequently move, but when motile, most cones within a cyst are unaligned and move abnormally (Noguchi et al., 2008). We found that across genetic backgrounds, Arp2L-generated actin cones were frequently unaligned, unlike Arp2s-generated cones (Fig. 3D-F, S3F). Trailing cones within a cyst often appear needle-like, yet most Arp2L-generated cones appear fan-shaped (Fig. 3D), unlike those observed with a complete loss of Arp2/3 activity (Noguchi et al., 2008). Thus, Arp2/3 appears active, but activity is suboptimal with the microexon present. We quantitatively confirmed that mRNA expression in the testis was comparable between the splice variants (Fig. S3I-J), as seen with overall adult fly expression (Fig. S3G-H). Due to a lack of an appropriate antibody to detect untagged Arp2, we cannot rule out definitively that Arp2s and Arp2L protein levels differ during sperm individualization; thus, it remains a possibility that the microexon may regulate Arp2 protein levels in specific cell types, potentially providing a mechanism to control Arp2/3 activity, analogous to a microexon found in a neuronal protein (Lin et al., 2020). Regardless of the mechanism, this cone defect reveals that the Arp2L microexon impacts Arp2/3 in certain tissue contexts.

Because actin polymerization drives cone motility (Noguchi and Miller, 2003), we also assessed Arp2L-generated cones for defects in movement. We cultured testes *ex vivo* and tracked cone movement live by expressing a GFP-tagged protein that localizes to the leading edge (Arp53D, Fig. 3G) (Schroeder et al., 2021). We found that Arp2s-generated cones moved at approximately 3 µm/min (Fig. 3G-H, Video 1), as expected for wildtype cones (Noguchi and Miller, 2003), yet Arp2L-generated cones would sometimes stall, followed by abrupt re-initiation of motility (Fig. 3G, Video 2). Thus, the overall speed at which Arp2L-generated cones traveled was slower (Fig. 3H), and the speed at which cones moved within a cyst varied more widely (Fig. 3I). We also observed that most cones exhibiting stalls were unaligned (Fig. 3J), yet the unaligned cones within a cyst moved synchronously (Fig. 3G). Thus, cones that appear to trail behind are not moving more slowly than other cones in a cyst; instead, they may trail due to a structural defect in the actin network. Currently, it is mechanistically unclear how actin branches coordinate alignment and proper motility.

These defects do not significantly reduce male fertility (Fig. S3K) or sperm production (Fig. S3L). Because males produce many sperm, partial defects in individualization usually have minor impacts on overall fertility (Stromberg et al., 2023). Nonetheless, these findings indicate that expressing Arp2L alone is suboptimal for individualization, which may explain why Arp2L tends to be less expressed than Arp2s in the testis (Fig. 1C). The cone defects serve as a readout for divergence of the splice variants *in vivo*, and additional phenotypes likely exist beyond the testis.

### The microexon sequence underlies the functional divergence of Arp2L in the testis

The actin cone defects found in males expressing only Arp2L may arise due to the sequence of the microexon or the mere lengthening of the D-loop by five residues. We tested whether the sequence led to the observed defects by replacing the microexon with five glycine residues. This strategy effectively “erases” the side chains but maintains the length and flexibility of the Arp2L D-loop (Fig. 4A). We found that our “5X-Gly” chimera fully rescued zygotic viability (Fig. 4B), and male flies displayed a lack of unaligned cones in the testis (Fig. 4C). Thus, the increased length of the D-loop alone does not underly defects in Arp2L-generated cones. We then tested whether cones were impacted by swapping the Arp2L microexon with the distinct, positively charged human Arp2 microexon (Fig. 4A). Adult flies were viable with the “human swap” chimera (Fig. 4D), and interestingly, the testis did not exhibit unaligned cones across genetic backgrounds (Fig. 4E, Fig. S3Q). All chimeras exhibited comparable mRNA expression in whole flies and the testis (Fig. S3J, M-P). In sum, the microexon sequence underlies the observed functional divergence of Arp2L *in vivo*, suggesting that the *Drosophila* Arp2L microexon may be maladaptive for individualization.

**Figure 4:**
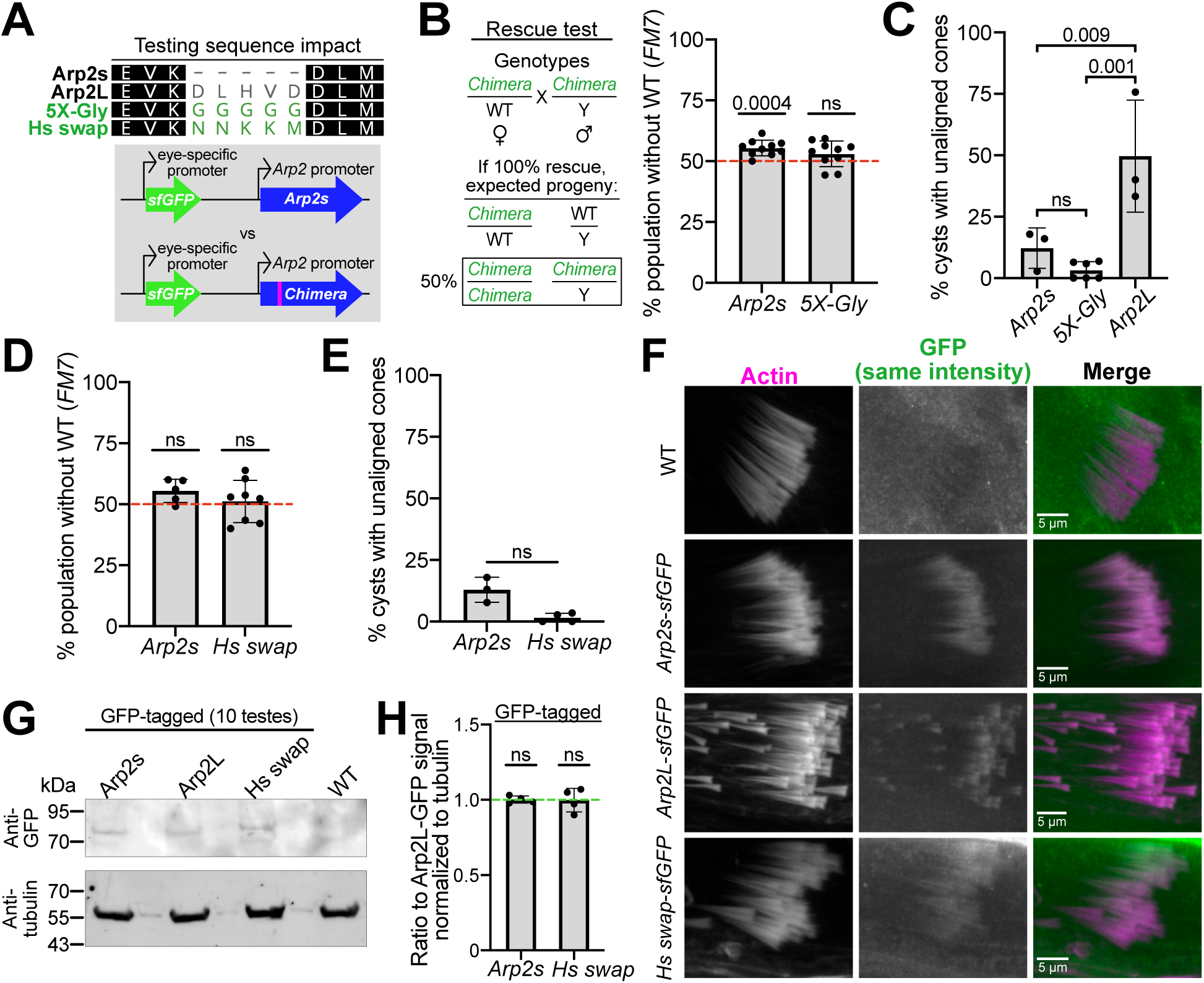
The sequence of the microexon underlies Arp2L functional divergence in the testis. **A)** Top, alignment of the *Drosophila* Arp2 microexon with the “5X-Gly” and “human swap” (or “Hs swap”) chimeras. Bottom, schematic of the transgenes encoding either *Arp2s* or the chimeras *5X-Gly*, or *Hs swap*, inserted into the *Arp2*-KO locus as in Fig. 3A. **B)** Left, schematic showing that females heterozygous for the *5X-Gly* chimera were crossed to hemizygous males to test for rescue of *Arp2*-KO lethality as in Fig. 3B. Right, quantification of percent of total progeny without the WT Arp2 allele (balancer *FM7*) and homozygous for the transgene. Dotted line indicates the expectation based on Mendelian inheritance. Statistical test: a one-sample ratio t-test with the hypothetical value being 50%. *5X-Gly* transgene fully rescues *Arp2*-KO zygotic lethality similar to *Arp2s*. Number of KO-*Arp2s* progeny was slightly higher than expected, likely indicating a small fitness cost from the *FM7* balancer. Average number of flies counted per replicate: *Arp2s*: 109, *5X-Gly*: 102 **C)** Quantification of the percent of cysts with unaligned cones in KO-*Arp2s*, KO-*Arp2L*, and KO-*5X Gly* males. Each datapoint represents a single cytological experiment quantifying total individualizing cysts across about 10 males per genotype. One datapoint for Arp2s is shared with panel E. Statistical test: one-way ANOVA with a Tukey’s multiple comparisons post-hoc test. 5X-Gly generated cones are significantly less unaligned than Arp2L-generated cones. **D)** Test for rescue of *Arp2*-KO lethality with Hs swap chimera, as in panel B. Average number of flies counted per replicate: *Arp2s*: 121, *Hs swap*: 45. **E)** KO-*Arp2s* cone alignment was compared to KO-*Hs swap* cones as in panel C. Statistical test: unpaired t-test with Welch’s correction, showing no difference. One datapoint for Arp2s is shared with panel C. **F)** Fluorescent images of actin cones in *w^1118^*males that encode the wildtype *Arp2* locus but also encode for *Arp2s, Arp2L* or Hs swap with a C-terminal sfGFP tag. Each transgene is under control of the Arp2 promoter and inserted into the second chromosome. Despite the presence of endogenous Arp2, all GFP-tagged proteins localize to the front of actin cones as expected for WT (no GFP observed in negative control). GFP was imaged with the same laser intensity and displayed with the same gray levels across samples. **G)** Western blot probing testis lysate from flies expressing GFP-tagged Arp2s, Arp2L, or Hs swap. Testes from flies not encoding a GFP-tagged protein (*w^1118^*) were used as the “WT” control. Samples were probed with an anti-GFP antibody, and tubulin was used as a loading control. **H)** Quantification of the western in G and three other similar westerns with independently acquired samples. Graph shows ratio of GFP intensity from Arp2s-GFP or Human swap-GFP testis samples to that of the Arp2L-GFP sample on the same blot. GFP band intensities were normalized to tubulin. Statistical test: a one-sample ratio t-test with the hypothetical value being 1. Similar GFP intensities suggest that altering the D-loop sequence does not impact protein expression of GFP-tagged Arp2.

We next assessed Arp2/3 localization to actin cones by comparing Arp2s, Arp2L, and the human swap chimera with a C-terminal GFP tag, all of which were expressed under the same Arp2 promoter used previously. We expressed them in a WT background, as the GFP tag can reduce viability in *Arp2*-KOs, and found that all three localize to cones (Fig. 4F). Interestingly, cones with localized Arp2L-GFP are significantly unaligned compared to cones with Arp2s- or Hs swap-GFP (Fig. S3R), consistent with untagged protein in the *Arp2*-KO background (Fig. 3E-F, Fig. 4E). We verified that protein expression is comparable among the splice variants (Fig. 4G-H), indicating that the microexon does not significantly alter GFP-tagged protein levels and that cone unalignment is likely due to differences in Arp2/3 activity.

### Expressing *Arp2s* alone is evolutionarily unfavorable

The apparent lack of defects in flies expressing only *Arp2s* raised the question as to why *Arp2L* even exists. Because evolution preserved the microexon for over 600 million years, Arp2L likely plays a critical role in a specific cell type. Given the challenge of uncovering where Arp2L is required, we opted to take an evolutionary approach and conduct a “population cage experiment” to test whether any fitness cost arises when flies express only *Arp2s*. In this experiment, we co-cultured flies expressing only *Arp2s* (KO-*Arp2s*) with WT flies and scored progeny genotypes after multiple generations using GFP eye fluorescence to identify the KO-*Arp2s* allele. If the *Arp2s* allele persists or even increases in frequency, then the WT allele, which encodes both *Arp2s* and *Arp2L*, provides no fitness advantage. Because the outcome of an evolutionary experiment can depend on the starting genetic background (Jerison et al., 2017), we conducted this experiment with two KO*-Arp2s* lines that had been independently isogenized in the fly line used as the WT fly: *w^1118^*, which encodes the wildtype *Arp2* locus. We used *w^1118^* flies produced from isogenization (the “siblings” lacking GFP) so that the WT and KO-*Arp2s* flies have similar genetic backgrounds. We also chose *w^1118^* flies as the WT fly because, like KO-*Arp2s* flies, they carry a mutation that results in a lack of eye pigmentation, facilitating detection of the GFP eye marker. Our starting population was composed of 50% KO-*Arp2s* flies and 50% *w^1118^* flies. With each generation, we collected 100 progeny randomly (50 females and 50 males without noting genotype) for the subsequent generation (Fig. 5A). After five generations, we found that the WT allele outcompeted the *Arp2s*-encoding allele (Fig. 5B). The average frequency reached approximately 75-95% across all four replicates, suggesting the reproducible rise in the WT allele is likely not due to genetic drift. Thus, our evolutionary experiments suggest that expressing *Arp2s* alone comes at a cost. We infer that Arp2L confers an overall fitness advantage, likely through a tissue-specific role, which explains the conservation of the microexon sequence across all fly species (Fig. S1B).

**Figure 5:**
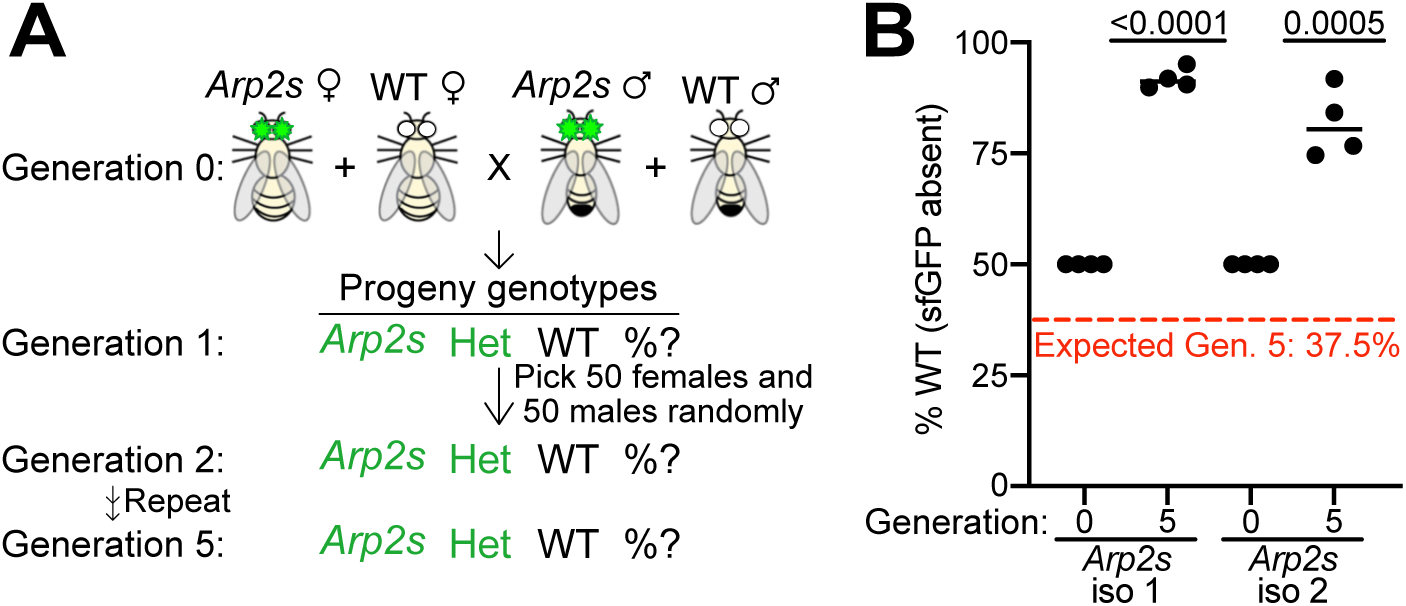
Expressing *Arp2s* alone is evolutionarily unfavorable. **A)** Schematic depicting the experimental setup of a population cage experiment. The starting population was composed of 50% KO-*Arp2s* females and males (isogenized in a *w^1118^* fly background) and 50% wildtype (WT) flies (*w^1118^*, same background as KO-*Arp2s* flies). The *Arp2s* transgene is genetically linked to a gene that encodes for sfGFP expression in the eye. For each passage, 100 progeny were randomly collected for each subsequent generation. Progeny are either homozygous for *Arp2s*, heterozygous (“Het”), or WT (lacking the *Arp2s* transgene). Generation five progeny were scored for sfGFP fluorescence in the eye. **B)** Quantification of the experiment in panel A, showing percent of the WT genotype. Two independently backcrossed KO-*Arp2s* fly lines were tested with four replicates each. Red line indicates the expected percentage of homozygous WT progeny if the WT allele does not confer a fitness advantage (37.5% based on the Hardy-Weinberg principle for an X-linked locus). The WT allele quickly outcompeted the *Arp2s* transgene. Statistical test: one-sample ratio t-test with the hypothetical value being 37.5%. Average population size per replicate: KO-*Arp2s* iso. 1: 562 flies; KO-*Arp2s* iso. 2: 394 flies.

### Concluding remarks

The context-dependent differences observed *in vitro* between Arp2s and Arp2L align with our findings that Arp2s and Arp2L are largely redundant *in vivo,* except in certain processes, such as late sperm development. The splice variants may functionally diverge in the presence of a tissue-specific regulator. Indeed, some microexons can alter tissue-specific protein interaction networks with significant functional consequences (Irimia et al., 2014; Li et al., 2015). Our work also demonstrates that the D-loop sequence impacts function, as observed with divergent Arp2 paralogs (Stromberg et al., 2023). Thus, the sequence variation of the microexon among species (Fig. 1B) may provide a means to modulate Arp2/3 function for unique biological contexts.

## Methods

### Fly husbandry and crosses

Fly lines were cultured at room temperature on yeast-cornmeal molasses-malt extract medium. Flies in all analyses were 1-5 days old. Flies referred to as “wildtype” are *w^1118^* flies (RRID:BDSC_3605) except for the analyses in Fig. 1C and Fig. S2, in which Oregon-R flies (RRID: DGGR_109612) were used. All crosses were set up with 5:2 females to males (unless otherwise noted), and vials with crosses were flipped every 3 days. For testing rescue of *Arp2*-KO zygotic lethality, at least five replicates were set up, and genotypes were scored per vial. For the male fertility assay, at least 7 replicate crosses were set up, and adult progeny were quantified no more than 20 days after the cross was established to avoid counting the subsequent generation.

### Generation of fly transgenics

The “KO-*Arp2s*,” “KO-*Arp2L*,” “5X-Gly,” and the “Hs swap” fly lines in this study were generated with the *Arp2*-KO fly line (balanced with *FM7*) from a previous study (Stromberg et al., 2023) to express *Arp2* transgenes in the absence of endogenous *Arp2*. The KO-*Arp2s* fly line was from a previous study (Stromberg et al., 2023), and the construct used to generate it was used to generate *Arp2L* via PCR mutagenesis. The 5X-Gly and human swap chimeras were also generated via PCR mutagenesis of the *Arp2s* construct. Mutagenesis was done with Phusion (NEB, Cat. No. M0530) as in a previous study (Stromberg et al., 2023). All tagless *Arp2* transgenes were flanked by the 1 kb upstream and downstream regions of the endogenous *Arp2* locus. For fly lines encoding *Arp2s* or *Arp2L* with a C-terminal superfolder GFP (sfGFP) tag (Pedelacq et al., 2006), the transgene was flanked by the 1 kb upstream region of the Arp2 locus and was Gibson cloned (NEB, Cat. No. E2611L) into a vector containing an *attB* site for site-directed transgenesis.

Constructs were sequence verified, most by whole plasmid sequencing, performed by Plasmidsaurus, which uses Oxford Nanopore Technology. Constructs were then midi-prepped (Takara Bio, Cat. No. 740410.5) and co-injected with PhiC31 (Bischof et al., 2007) into *attP*-encoding flies for site-directed transgenesis by Rainbow Transgenic Flies, Inc. For integration into the *Arp2*-KO locus on the X chromosome, tagless transgenes were injected into our *Arp2*-KO fly line, which encodes an *attP* site in the *Arp2*-KO locus, and G0 adults were crossed to the *Arp2*-KO fly stock to identify transformants and balance the transgenes with *FM7* (identified by a distinct eye shape). The *Arp2*-KO locus was identified by DsRed fluorescence in the eye, and *Arp2* transgenes were identified by GFP eye fluorescence because the transgenes are closely linked to a gene encoding sfGFP under the control of an eye-specific promoter (3XP3). For backcrossing, flies were backcrossed to *w^1118^* for five generations before balancing with *FM7*. To test for phenotype consistency across genetic backgrounds, tagless Arp2s, Arp2L, and the human swap chimera were also inserted into the second chromosome, using stock BL9736 (Bloomington *Drosophila* Stock Center, RRID:BDSC_9736). We then crossed G0 adults to stock BL9736 to select for transformants. Second chromosome transgenes were subsequently balanced with *CyO* using a double-balanced fly line (*w^−^; CyO/Sp; Sb/TM6B)*. Once *Arp2*/CyO flies were obtained, they were crossed to *Arp2*-KO/*FM7* flies to generate a balanced stock with the *Arp2* transgenes in the *Arp2*-KO background (ex: *Arp2-KO/FM7; Arp2L/CyO*).

For flies encoding GFP-tagged Arp2, constructs for transgenesis were injected into stock BL9736 for incorporation into an *attP* site in the second chromosome of a wildtype fly line. G0 adults were crossed to BL9736 flies, and progeny with GFP fluorescence in the eye were selected, as GFP under the control of an eye-specific promoter (3XP3) was encoded in the original construct for transgenesis. All transgenics were sequence verified by extracting genomic DNA, as in a published protocol (Huang et al., 2009), and conducting a PCR (Phusion, NEB, Cat. No. M0530) using primers in Table S2. PCR products were gel-purified (Zymo) and sequenced by Plasmidsaurus.

### Population cage experiment

To test for a fitness cost in the absence of *Arp2L*, we performed a population cage experiment similar to a past study (Schroeder et al., 2021). For this assay, two different KO-*Arp2s* founder lines were independently backcrossed to a *w^1118^* fly line for five generations. The *w^1118^* progeny (lacking eye fluorescence) from backcrosses with the *Arp2*-KO background were selected and amplified for use as the “wildtype” fly in the population cage experiment, as this *w^1118^* fly has the closest genetic background to the *w^1118^*-isogenized KO-*Arp2s* fly lines. The *Arp2s* allele was identified by GFP fluorescence in the eye, and the closely linked KO allele was identified by DsRed fluorescence in the eye. Virgin female and male KO-*Arp2s* and *w^1118^* flies (25% of the population per sex and genotype) were combined into a bottle. Four replicate bottles, or “cages,” were created for each independently isogenized KO-*Arp2s* line. The four crosses for each KO-*Arp2s* line were passaged every two weeks at 25°C. The parents of each generation were allowed to lay for 1 week and then removed and frozen for quantifying later. After progeny hatched, 100 were randomly selected (without the use of fluorescence) for the subsequent generation. The remaining progeny were frozen for quantifying genotypes via GFP fluorescence at a later time, along with the parents of the subsequent generation. Progeny from the fifth generation were scored and classified as either GFP negative (only WT allele present) or GFP positive (heterozygous or homozygous).

### Expression analyses

Whole flies or fly tissue were dissected and flash frozen. At least 50 embryos, 20 larvae, 20 pupae, and 10-20 adult fly tissues were pooled for RNA extractions and all expression analyses. For testis dissections, 40 testes were collected. Samples were thawed and homogenized in TRIzol (Fisher Scientific, Cat. No. 15-596-026), and RNA was extracted as previously done (Stromberg et al., 2023). Purified RNA was then used to generate cDNA using SuperScript III First-Strand Synthesis (Thermo Fisher Cat. No. 18080051) using oligo dTs as the primer, per manufacturer’s instructions (Invitrogen). For each sample, a “-RT” control was performed in which the reverse transcriptase was replaced by water; these samples were used for PCR reactions with the primers targeting *Arp2* transgenes to determine if contaminating genomic DNA was present. Oregon-R cDNA was used to conduct quantitative RT-PCR (RT-qPCR, see below). The cDNA from transgenic flies was used to conduct RT-qPCR or RT-PCRs with Phusion (NEB, Cat. No. M0530) and analyzed by gel electrophoresis. For RT-PCRs, 25 cycles were conducted (unless otherwise noted), and reactions were loaded onto a 1% agarose gel for visualization.

For RT-qPCRs, iTaq Universal SYBR Green Supermix (BioRad, Cat. No. 1725121) was used with approximately 50 ng of cDNA in each reaction. The manufacturer’s instructions were followed for the Applied Biosystems 7500 Real-Time PCR System (Life Technologies). A standard curve method was conducted with the following parameters: 15 s denaturation at 95°C, 60 s extension time at 60°C with 40 cycles. For assessing *Arp2s* versus *Arp2L* expression in wildtype (Oregon-R) flies, primers targeting *Arp2s* and *Arp2L* aligned at the microexon junctions and were picked based on single peaks in the melt curve analysis and demonstrated specificity. Specificity of primers was evaluated by using cDNA from KO-*Arp2s* and KO-*Arp2L* flies as the template. We selected *Arp2L*-targeted primers that successfully detected *Arp2* from KO-*Arp2L* flies (low C_t_), while failing to amplify Arp2s from the KO-*Arp2s* fly line (high C_t_ or undetermined C_t_). The same strategy was used to select *Arp2s*-targeting primers, using KO-*Arp2L* primers as a negative control. We found that primer pairs had comparable amplification efficiencies, evaluated by a dilution series of Oregon-R genomic DNA. Replicates were performed with the cDNA diluted fresh each time to control for variation in pipetting._For quantitatively assessing *Arp2* transgene expression in the testis across transgenics, the same set of Arp2 primers, which aligned downstream of the microexon, were used for RT-qPCRs and are listed in Table S2. Forty testes were dissected for each biological replicate (three total replicates). All primers and raw C_t_ values are listed in Table S2, and all C_t_ values were normalized to those for *rp49*.

### Tissue Dissections and Immunofluorescence

Virgin flies were aged approximately 3-4 days before dissecting. Females were fed yeast paste at least 24 h before dissecting the ovaries to promote oogenesis. For immunofluorescence, ovaries and testes were dissected in PBS and then fixed in 4% paraformaldehyde as done previously (Stromberg et al., 2023). For actin staining, tissues were incubated with 2 µM SiR-Actin (Cytoskeleton, Inc, Cat. No. CY-SC001) for 2-3 h at room temperature or overnight at 4°C. To visualize GFP-tagged Arp2s and Arp2L, fixed testes were permeabilized with 0.3% Triton X-100 for 30 min, followed by several washes with PBS, 0.1% Triton (PBST). Testes were then incubated overnight at 4°C with anti-GFP (Abcam ab13970, RRID:AB_300798) at 1:500 in PBST with 3% bovine serum albumin (BSA, Fisher Scientific, Cat. No. BP1605-100). After washing testes several times, they were incubated for 2 h at room temperature with anti-chicken Alexa Fluor Plus 488 (Thermo Scientific, Cat. No. A32931, RRID:AB_2762843) at 1:2,500 and 2 µM SiR-Actin (Cytoskeleton, Inc., Cat. No. CY-SC001). All tissues were probed for DNA with Hoechst 33342 (Invitrogen, Cat. No. H3570) for 10 min at room temperature before mounting on slides with ProLong Diamond Antifade Mountant (Thermo Fisher, Cat. No. P36970).

### Microscopy and Image Analysis

#### Analyzing whole flies

Markers for transgenes were identified using a Leica M165 FC stereomicroscope equipped with filters to allow for visualization of eye fluorescence (GFP and DsRed). To observe bristles, flies were anesthetized, examined using brightfield with the Leica M165 FC stereomicroscope, and imaged with a Leica Flexacam C5 camera.

#### Fixed images

All fluorescent samples were imaged using an inverted Ti2 Nikon AX-R confocal microscope and NIS-Elements (Nikon, RRID:SCR_014329). Images were subsequently analyzed using Fiji, RRID:SCR_002285 (Schindelin et al., 2012; Schneider et al., 2012), and cones were classified as misaligned if five or more were trailing behind the remaining cones within a cyst. For actin cone images in Fig. 3, Nikon’s denoise.ai algorithm was used to remove background. For measuring the area of seminal vesicles, seminal vesicles attached to the testis, were stained with SiR-Actin and Hoechst, and the line tool in Fiji (Schindelin et al., 2012; Schneider et al., 2012) was used to outline the entire organ.

#### Live imaging of actin cones

For imaging cone motility, KO-*Arp2s* and KO-*Arp2L* flies were crossed to flies encoding GFP-Arp53D (Schroeder et al., 2021). Testes from the male progeny of the cross were dissected in a dish containing Schneider’s *Drosophila* media (Fisher Scientific) with 10% FBS (Sigma, Cat. No. 12306C). Then 1-2 testes were placed in a glass-bottom microwell dish (MatTek, Cat. No. P35G-1.5-10-C) and submerged in 50-100 µL of media. Images were then acquired on an inverted Ti2 Nikon AX-R confocal microscope over 1-2 hours. Z-stacks of GFP fluorescence were acquired every 6 min, as done in a previous study that imaged cones live (Noguchi and Miller, 2003). Maximum projections were then made for each time point in Fiji, and the distance cones moved from frame to frame was measured using the line tool. In KO-*Arp2L* males, even misaligned cones moved synchronously, allowing for tracking the front of the group of cones from frame to frame. To calculate overall speed, the distance traveled in each frame was summed and divided by the total time traveled (6 min per frame). Multiple individualizing cysts were measured within a testis, and six to seven males were imaged per genotype. Only cones that were motile for at least one frame of the acquisition were quantified to ensure that the analyzed cones had initiated motility.

#### GFP-tagged Arp2 confocal and immunoblot images

All cytological samples of testes expressing GFP-tagged Arp2s, Arp2L, or the human swap chimera were imaged with the same laser intensity for actin (Cy5 channel) and GFP (488 channel). Images in Figure 4 were then made by generating maximum projections in Fiji, RRID:SCR_002285 (Schindelin et al., 2012; Schneider et al., 2012), with 25 slices total, and the gray levels were kept constant across images. Gray levels for actin were not kept constant as actin staining showed variability across slides from different experiments. For quantification of western blot images, Fiji (Schindelin et al., 2012; Schneider et al., 2012) was used to measure the raw integrated density of GFP and tubulin bands using the same sized region of interest across samples. Each GFP band measurement was divided by the measurement for the corresponding tubulin band. Normalized intensities for Arp2s-GFP and human swap-GFP were divided by that of the Arp2L-GFP signal to compare levels relative to Arp2L-GFP.

### Protein gels and Immunoblots

All protein samples were denatured with 4X NuPAGE LDS sample buffer (Thermo Fisher, Cat. No. NP0007), boiled at 95°C for 5 min, and loaded onto NuPAGE 4-12% Bis-Tris gels (Thermo Fisher, Cat. No. NP0321BOX). Gels were run with MES SDS buffer (Thermo Fisher, Cat. No. NP0002). Gelcode Blue Safe Protein Stain (Fisher Scientific, Cat. No. PI24594) was used to visualize purified proteins following the manufacturer’s instructions. For assessing GFP-tagged Arp2 protein expression, a western blot was carried out. Proteins were transferred to a PVDF membrane then incubated with 5% milk in Tris-buffered saline (TBS, Sigma Cat. No. T5912). Membranes were probed with primary antibody overnight at 4°C or at room temperature for 1 hr, followed by three washes with TBS. Then secondary antibodies (anti-chicken Alexa Fluor 647, Invitrogen RRID: AB_2535866; anti-mouse Alexa Fluor 568, Thermo Fisher, RRID: AB_2534072) were diluted to 1:1,000, applied to the membrane and incubated for 1 hr. Immunoblots were then imaged with 647 nm and 546 nm using a ChemiDoc MP Imaging System (BioRad). Primary antibodies were anti-GFP (chicken, 1:500, Abcam, ab13970, RRID: AB_300798) and anti-tubulin (DM1A, Thermo Scientific, 1:1,000, Cat. No. 62204).

### Analysis of Embryonic Development

To assess embryo hatching, crosses were set up in a cage fitted with a 35 mm plate containing grape agar (Genesee Scientific) and a drop of yeast paste. Crosses included the same ratio of females to males across all genotypes tested in each experiment (generally 10:2 females to males). Flies were acclimated to the cage for approximately 24 hours. Then the plate was switched out at the end of the day, and the embryos that were laid overnight were transferred to a fresh grape-agar plate. The number of hatched embryos (identified by a deflated morphology) were quantified 24 and 48 hours following the transfer to a new plate. Little additional hatching occurred beyond this timepoint.

To assess embryos for fertilization and the stage of development, embryos were washed with PBS and dechorionated with freshly diluted 50% bleach for 30-60 s. Then the embryos were washed and placed into a mounting chamber, described in detail previously (Figard and Sokac, 2011). Briefly, two pieces of double-sided stick tape were adhered to a slide with about 3-4 mm between them. Dechorionated embryos were placed between the pieces of tape and then halocarbon oil 700 (Sigma Aldrich, Cat. No. 9002-83-9) was pipetted on top of them to prevent dehydration. A coverslip was placed on top, and the embryos were imaged using brightfield on an inverted Ti2 Nikon AX-R confocal microscope. If cells were apparent in the embryos, they were classified as fertilized. Embryos were also classified as “late embryogenesis” if invaginations were observed (Figard and Sokac, 2011).

### Analysis of Actin rings and Thoracic Bristles

For quantifying the diameter of actin rings in ovaries, only stage 10 egg chambers were imaged and only rings that were perpendicular to the imaging plane were measured. Z-stacks were acquired and the slice in the middle of the ring was used for quantification. Using NIS-Elements (Nikon, RRID:SCR_014329), a line was drawn across the ring, with endpoints just outside the points of actin fluorescence. Only structures that were confidently identified as rings were quantified; sometimes the actin probe (SiR-Actin, Cytoskeleton, Inc. Cat. No. CY-SC001) appeared to stain lipid globules, which were avoided in quantification.

Adult virgin flies were scored for bristle defects soon after eclosion and then reassessed later in the day. Bristle defects were identified as observed in past studies that found bristles lacking Arp2/3 activity exhibit kinks or severe bends in macrochaetes and microchaetes (Frank et al., 2006; Koch et al., 2012).

### Sequence and Structural Analyses

All sequence alignments were done using MUSCLE, RRID:SCR_011812 (Edgar, 2004), in the Geneious software (RRID:SCR_010519) and visualized with Geneious (Kearse et al., 2012) or Jalview, RRID:SCR_006459 (Waterhouse et al., 2009). Species trees were obtained using the TimeTree database, RRID:SCR_021162 (Kumar et al., 2022), and visualized in Geneious (Kearse et al., 2012). *Arp2L* sequences were obtained across diverse species by conducting a tBLASTn search (Camacho et al., 2009), RRID:SCR_011822. The query was the human Arp2L sequence for vertebrates and the *Drosophila melanogaster* Arp2L sequence for invertebrates. Given NCBI predicted some Arp2L sequences based on genomic sequence, RNA databases in the NCBI Sequence Read Archive (RRID:SCR_004891, Table S1) were further surveyed to determine if unusual invertebrate microexons were indeed expressed. tBLASTn, using the corresponding species’ Arp2L sequence, was conducted with the RNA databases for a subset of species (Table S1). If the microexon was identified in the tBLASTn hits, then it was used in the sequence alignment in Fig. 1B. For structural analyses, AlphaFold 3 (Abramson et al., 2024) was used to predict the structure of the *Drosophila melanogaster* Arp2/3 complex, containing Arp2s or Arp2L with Arpc3A or Arpc3B. All structure predictions included two actin monomers, four ATP molecules, and four Mg^2+^ ions as ligands, which bind actin, Arp2, and Arp3. All structures, including electrostatic potential, were visualized using ChimeraX (Meng et al., 2023). For mapping sequence conservation on Arp2/3 structures, sequence alignments were used with the “Render by Attribute” tool in ChimeraX, RRID:SCR_004097 (Meng et al., 2023).

### Recombinant protein expression and purification

Recombinant *Drosophila* Arp2/3 was cloned using the BigBac system (Weissmann et al., 2016) as was done for human Arp2/3 (Zimmet et al., 2020). All subunits were codon-optimized (Genewiz), and a 1X FLAG-tag was encoded onto the C-terminus of Arpc3A/B, similar to the purification strategy for the human recombinant Arp2/3 (Zimmet et al., 2020). The final pBig2abc construct (RRID:Addgene_80617) encoding all seven subunits of the Arp2/3 complex was fully sequenced (Plasmidsaurus) to ensure a lack of mutations and then transformed into Max Efficiency DH10Bac competent cells (Thermo Fisher). Positive colonies were selected using blue-white screening (Weissmann et al., 2016). The bacmid was isolated from positive (white) colonies using the Wizard Plus SV Minipreps DNA purification system (Promega). The pure bacmid was then immediately used to transfect Sf9 cells (Thermo Scientific Cat. No. A35243). The ExpiSf9 baculovirus expression system (Thermo Fisher, Cat. No. A3767806) was used for baculovirus generation and protein expression. To generate baculovirus, ExpiFectamine Sf transfection reagent (Thermo Scientific, Cat. No. A38915) was used to transfect ExpiSf9 cells with purified bacmid (per manufacturer’s instructions). The subsequent P0 virus was isolated approximately 5 days following transfection by spinning down the cells and storing the virus-containing supernatant at 4°C. The P0 virus was used to infect ExpiSf9 cells and generate P1. The P1 virus was finally used to infect 400 mL of 5 million cells/mL for protein expression. The infection took place for 3 days before spinning down the cells and flash freezing the pellet for affinity purification.

An approximately 5 g pellet of Sf9 cells (Thermo Scientific Cat. No. A35243) was used to purify 2-4 mg of Arp2/3 protein. To lyse the cells, the pellet was quickly thawed and resuspended in 20 mM HEPES (pH 8), 200 mM NaCl, 1% Triton X-100, 2 mM TCEP, 2 mM PMSF, and cOmplete EDTA-free protease inhibitor cocktail (Millipore Sigma, Cat. No. 11836170001). Following Dounce homogenization, the lysate was spun at 20,000 rpm (rotor JA 25.50) for 30 minutes at 4°C. About 1 mL of anti-FLAG resin (GenScript, L00432) was washed using lysis buffer and then incubated with the clarified cell lysate for 1-2 h at 4°C. The resin was then transferred to a gravity flow column and washed with buffer containing 20 mM HEPES (pH 8), 200 mM NaCl, and 2 mM TCEP. To elute, the resin was incubated with wash buffer containing 0.3 mg/mL 3X FLAG peptide (Sigma, Cat. No. F4799) for 30 minutes. Approximately eight 1-mL fractions were collected. Fractions containing the most protein were then combined and dialyzed with buffer containing 20 mM HEPES (pH 8), 100 mM NaCl, and 1 mM TCEP using dialysis tubing with a molecular weight cutoff of 12-14 kDa (Cole-Parmer, Cat. No. EW-02900-02). Following overnight dialysis, the protein was concentrated using an Amicon ultra-15 centrifugal filter unit (Millipore Sigma, UFC901008) to about 20 µM, then aliquoted and flash frozen. For intact mass spectrometry, purified Arp2/3 complexes were analyzed by LC/MS using a Sciex X500B QTOF mass spectrometer coupled to an Agilent 1290 Infinity II HPLC. The acquired mass spectra for the proteins were deconvoluted using BioPharmaView v. 3.0.1 to obtain the molecular weights.

For purification of the VCA domain of *D. melanogaster* WASP, a similar approach was used as was done for purifying human WASP (Cory et al., 2003; Yarar et al., 2002). The VCA domain of *Drosophila* WASP (aa 421-527) was codon optimized for bacterial expression and cloned into the bacterial expression vector pGEX-4T-1 (empty backbone of RRID:Addgene_73375). Following sequence verification (Plasmidsaurus), the construct was transformed into *Escherichia coli* BL21 DE3 cells (NEB, Cat. No. C25271). Cells were grown to an OD_600_ of 0.6-0.8 and induced with 0.4 mM isopropyl β-d-thiogalactoside at 37°C for 3 h. Cells were spun down and flash frozen. For affinity purification, a ∼5 g pellet was resuspended in lysis buffer containing 20 mM HEPES (pH 7), 300 mM KCl, 0.01% NP-40, 1 mM TCEP, 1 mM PMSF, and cOmplete EDTA-free protease inhibitor cocktail (Millipore Sigma, Cat. No. 11836170001). An Emulsiflex was used to lyse the cells. Clarified lysate was then incubated with 1 mL of glutathione Sepharose 4B resin (Cytvia, Cat. No. 17075601) at 4°C for 1 hour. The batch resin was gently spun, and the supernatant was removed. The resin was then resuspended in 10 mL of wash buffer containing 50 mM HEPES (pH 7), 300 mM KCl, 1 mM TCEP and added to a gravity flow column. The resin was washed with at least 10 mL of buffer, and the protein was then eluted with wash buffer including 10 mM glutathione (Millipore Sigma). The fractions with the largest amount of protein were then gel filtered with an SEC650 column (BioRad, Cat. No. 7801650) equilibrated with buffer containing 20 mM HEPES pH 7, 50 mM KCl, 1 mM MgCl_2_, 1 mM EGTA, and 0.5 mM TCEP. The protein was subsequently flash frozen at approximately 20 µM.

### Actin polymerization assays

Rabbit skeletal muscle actin (1-2 mg, Cytoskeleton Inc., Cat. No. AKL99) and pyrene-labeled actin (1 mg, Cytoskeleton, Inc., Cat. No. AP05) were separately resuspended in approximately 300 µL of G-buffer (2.0 mM Tris-Cl (pH 8), 0.2 mM ATP, 0.1 mM CaCl_2_, 0.2 mM TCEP) and then dialyzed for 2 days, exchanging with fresh buffer about 3 times. Actin was then spun at 100,000 x g for 2 h at 4°C, and to avoid a filamentous actin pellet, 80% of the supernatant was used to isolate only monomeric actin for subsequent gel filtration. Actin was gel filtered using an SEC650 column (BioRad, Cat. No. 7801650) equilibrated with G-buffer. The fractions containing actin were then returned to dialysis tubing in G-buffer at 4°C for use in polymerization assays the following day.

Polymerization assays were conducted as in a previous study (Balzer et al., 2020). Unlabeled and pyrene-labeled actin were combined at a final concentration of 10 µM, with 10% pyrene-labeled and 1 mg/mL BSA (Fisher Scientific, Cat. No. BP1605-100) to prevent nonspecific sticking. For the polymerization reaction, actin was diluted to a final concentration of 2 µM. A glass-bottom 96-well half-area microplate (Corning 3880) was set up with actin in one well and the neighboring well containing test proteins, either Arp2/3 or Arp2/3 and GST-VCA. Proteins were each diluted in 1X KMEI (10 mM Imidazole pH 7, 50 mM KCl, 1 mM EGTA, and 1 mM MgCl_2_) immediately before addition to the test-protein solution. Upon addition to the well containing actin, the test-protein solution consisted of a final concentration of 1X KMEI, 0.2 mM ATP, and 1 mM TCEP. Arp2/3 and GST-VCA were at 20 nM and 100 nM, respectively, for all assays. For assays including CK-666 (Sigma, Cat. No. SML0006) or CK-869 (Sigma, Cat. No. C9124), the drug was included in the well containing test proteins and diluted to a final concentration of 100 or 200 µM in the test reaction. The control for Arp2/3 inhibitors contained a volume of DMSO (Sigma, Cat. No. D2653) equivalent to that used for the drugs. Immediately upon addition of the test-protein solutions to actin, fluorescence readings were collected every 15 s for 45 min using a Tecan Spark microplate reader with the software SparkControl. Settings included an excitation wavelength of 365 nm with an excitation bandwidth of 10 nM, an emission wavelength of 407 nm with an emission bandwidth of 20 nm, a gain of 80, 30 flashes, and 40 µs of integration time.

### Statistical Analyses and Graphs

All graphs were made in GraphPad Prism (version 10, RRID:SCR_002798), and statistical significance was evaluated using the statistical tools in GraphPad Prism, noted in all figure legends. Values for experimental replicates are displayed on all graphs when possible. For most analyses comparing two different samples, an unpaired, two-tail t-test with Welch’s correction was conducted. When comparing more than two samples, a one-way ANOVA with a Tukey’s multiple comparisons post-hoc test was done, comparing the means of each sample. For experiments with Arp2/3 inhibitors, actin polymerization data were analyzed with a two-way ANOVA, with protein samples and inhibitor treatments as factors; a Fisher’s LSD post hoc test was used for prespecified pairwise comparisons. To test the statistical significance of a ratio from the expected ratio, a one-sample ratio t-test was conducted, testing significance against a hypothetical value, noted in the legends.

For actin cone analyses, cytology was done at least three times. Each cytology experiment included approximately 10 males, and all individualizing cysts across all testes were scored for unaligned versus aligned cones, with the percent of unaligned cones within that population graphed per replicate. Approximately 30 individualizing cysts were scored per genotype per replicate.

To compare relative actin polymerization rates, a similar approach was taken as in a previous study (Zimmet et al., 2020). Using GraphPad Prism (RRID:SCR_002798), the first derivative of each polymerization curve was taken. The maximum first derivative of each curve was normalized by dividing it by the maximum first derivative for actin alone, measured the same day as the experimental samples. To display the polymerization of actin over time, all curves were baseline-corrected by subtracting the average of the first three values.

## Data Availability

Most data are made transparent in the graphs and provided in the supplemental figures and tables. An extensive compilation of raw data will be made available in the UT Southwestern Research Data Repository, and any other data are available from the corresponding author upon request.

## Supplemental Material Summary

Supplemental materials include three figures, two tables, and two videos. Fig. S1 contains an expanded alignment of Arp2L sequences shown in Fig. 1B and a schematic depicting the strategy for generating transgenic flies in Figures 3-4. Fig. S2 contains additional supporting data for the *in vitro* results in Figure 2 and includes an analysis of embryonic development, showing differences between genetic backgrounds when flies express only *Arp2L*. Fig. S3 contains additional analyses of tissues, fertility, cones, and transgene expression levels, related to Figures 3-4. Table S1 lists the NCBI databases used to verify expression of the Arp2L microexon in invertebrates, related to Fig. 1B. Table S2 includes the primers used for sequence verification of transgenic fly lines and for RT-qPCRs, related to Fig. 1C. Table S2 also lists the raw cycle threshold values from all RT-qPCRs and the results of intact mass spectrometry analyses, related to Fig. 2B. Videos 1 and 2 are related to Fig. 3G, showing Arp2s- and Arp2L-generated cone motility.

## Video Legends

**Video 1: Arp2s-generated cone motility, related to Figure 3G**

Testis from KO-*Arp2s* fly cultured *ex vivo*. Testis also expresses GFP-tagged Arp53D, which localizes to the leading edge of cones. GFP-Arp53D at cones was imaged with the 488 channel, and a z-stack was acquired every 6 minutes. A maximum projection of slices acquired over 108 min of imaging time is shown at one frame per second. Scale bar 20 µm. The corresponding montage in Figure 3G is rotated 90 degrees.

**Video 2: Arp2L-generated cone motility, related to Figure 3G**

Testis from KO-*Arp2L* fly cultured *ex vivo*. Testis also expresses GFP-tagged Arp53D, which localizes to the leading edge of cones. GFP-Arp53D at cones was imaged with the 488 channel, and a z-stack was acquired every 6 minutes. A maximum projection of slices acquired over 108 min of imaging time is shown at one frame per second. Scale bar 20 µm.

## Supplemental Tables

**Table S1: Expression Databases**

The NCBI Sequence Read Archive (SRA) databases used to verify microexon expression are listed for invertebrates shown in Fig. 1B.

**Table S2: Primer sequences, raw values for RT-qPCRs, and intact mass spectrometry**

Intact mass spectrometry analyses of purified recombinant Arp2/3 complexes are provided. The primers used for sequence verification of transgenic fly lines and for RT-qPCRs are listed as well as raw cycle threshold (C_t_) values obtained from all RT-qPCR analyses. C_t_ values are also provided for experiments in which primers are tested for specificity. For example, primers targeting *Arp2L* were paired with cDNA from KO-*Arp2s* flies versus KO-*Arp2L* flies. *Arp2L*-targeting primers should have C_t_ values greater than 30 or undetermined when paired with cDNA from KO-*Arp2s* flies. Indeed, Arp2L-targeting primers only indicated true signal (C_t_ values less than 30) when paired with cDNA from KO-*Arp2L* flies. The same finding held true for *Arp2s*-targeting primers.

## Acknowledgments

We thank Michael Buszczak, David Corey, Miriam Osterfield, Andrew Lyon, and all Schroeder lab members for providing helpful feedback on the manuscript and experimental suggestions. We are grateful to Brad Nolen, Sofia Carlson, and Meghan Bacher for experimental advice on actin polymerization assays. We also thank Michael Rosen for the use of his fluorimeter, Michael Reese and Xuelian Luo for protein expression vectors, and the UTSW Proteomics Core for intact mass spectrometry analyses. Structural analyses and graphics were performed with UCSF ChimeraX (Meng et al., 2023), developed by the Resource for Biocomputing, Visualization, and Informatics at UC, San Francisco (supported by NIH R01-GM129325 and the Office of Cyber Infrastructure and Computational Biology, National Institute of Allergy and Infectious Diseases). This work was funded by the Cancer Prevention and Research Institute of Texas (RR210048), an NIGMS R00 (GM137038), an NIGMS R35 (GM156807), the UT Southwestern Endowed Scholars Program in Medical Science, and a UT System Rising STARs award (all funds to C.M.S.).

## Author Contributions

J.P.: data acquisition, data analysis, data curation, validation, generation of fly transgenics, writing, editing, and figure production. M.S.P.: data acquisition, data analysis, data curation, validation, generation of fly transgenics, editing, and figure production. C.M.S.: conceptualization, project management, funding acquisition, methodology, data acquisition, data analysis, data curation, validation, writing, editing, and figure production.

## Declaration of interests

The authors declare no competing interests.

---

**Figure S1:**
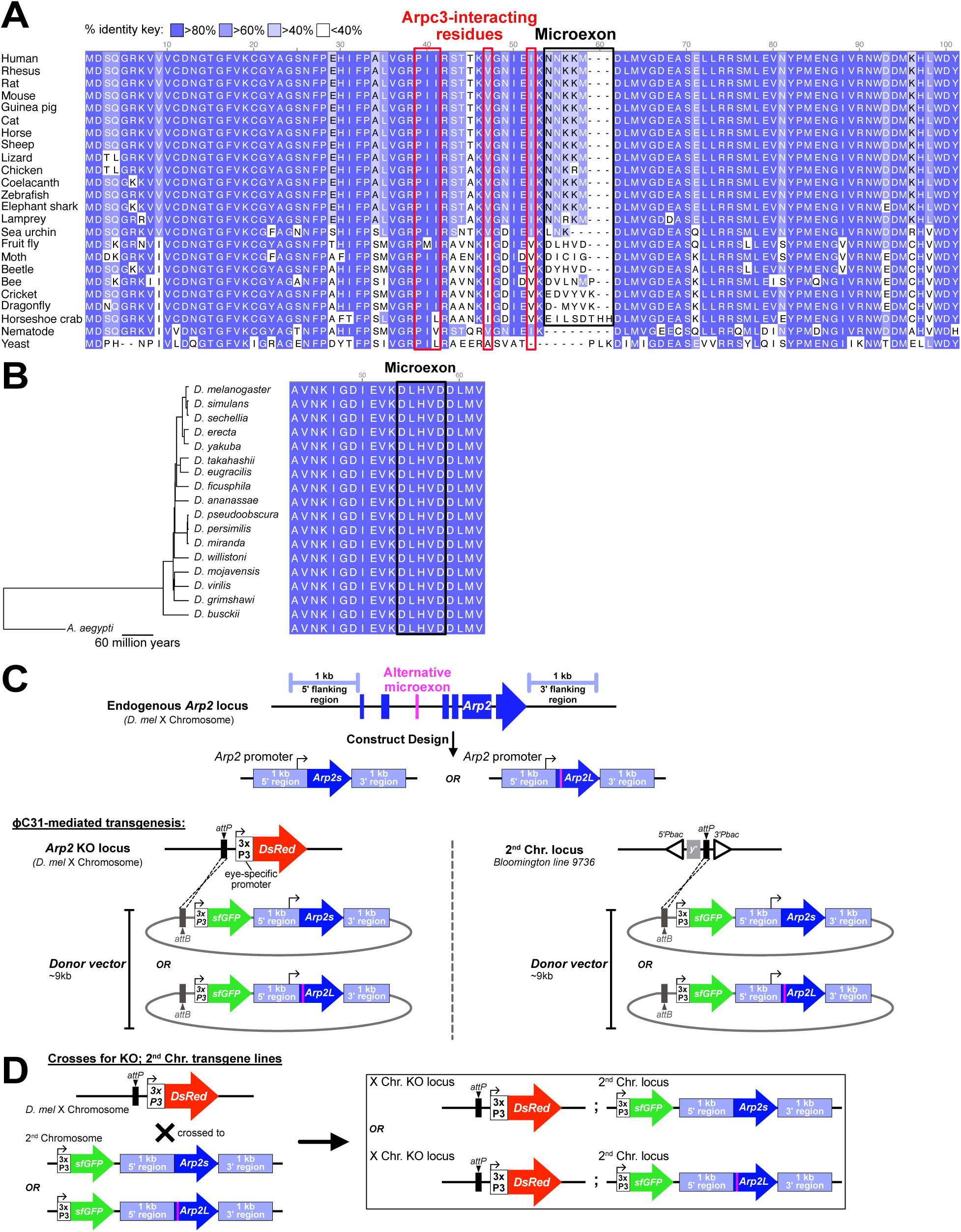
The Arp2 microexon diverges across 700 million years of eukaryotic evolution but is conserved in *Drosophila*, related to Figure 1. **A)** Protein sequence alignment including Arp2L orthologs from all species in Fig. 1B. The microexon and Arpc3-interacting residues (Ding et al., 2022) are boxed black and red, respectively. Key indicates the percent identity that corresponds to each color. **B)** Alignment of the Arp2L microexon (boxed in black) from 17 *Drosophila* species. Left, species tree with *Aedes aegypti* (mosquito) as the outgroup. The microexon is 100% conserved across Drosophilids. **C)** Schematic of the *Arp2s* and *Arp2L* transgene design. Endogenous *Arp2* contains six introns and is annotated on the minus strand of the *D. melanogaster* genome. For clarity, the endogenous locus is displayed in the 5′ to 3′ transcriptional orientation of Arp2. One-kilobase genomic fragments immediately flanking 5′ and 3′ of *Arp2*, including the native 5′ and 3′ UTRs and promoter, were cloned flanking intronless *Arp2s* and *Arp2L* coding sequences in the native transcriptional orientation. The resulting transgene cassettes were inserted into an *attB*-containing vector for PhiC31-mediated transgenesis. The vector also encodes an eye-specific *3XP3*-*sfGFP* marker, enabling identification of transgene-bearing flies by eye fluorescence. Vectors were integrated into the *attP* site in the *Arp2*-KO locus or an *attP* site on the second chromosome of an otherwise WT fly, the BL9736 fly line (Venken et al., 2006). **D)** Schematic summarizing the strategy of crossing wildtype flies encoding *Arp2s* or *Arp2L* on the second chromosome into the *Arp2*-KO background (identified by DsRed-positive eyes). Intermediate crosses to homozygose the *Arp2*-KO locus and *Arp2s/L*-encoding locus are not depicted. Transgenic lines were first balanced with *FM7* (*X* chromosome) and *CyO* (2^nd^ chromosome) and then crossed to select for homozygous *Arp2*-KO flies that lacked one or both balancers and encoded one or two copies, respectively, of *Arp2s* or *Arp2L*.

**Figure S2:**
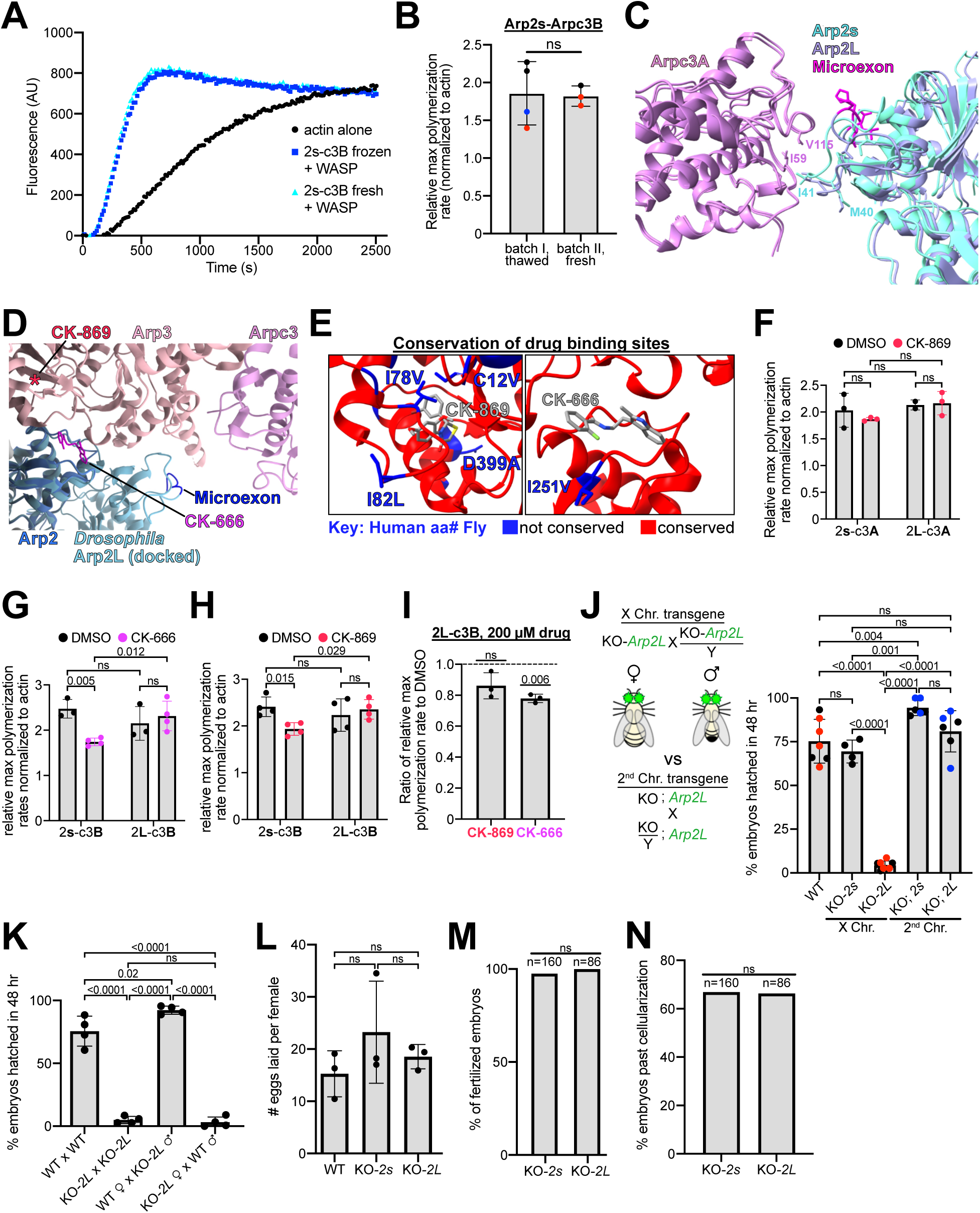
Additional *in vitro* data and analysis of an Arp2L phenotype inconsistent across genetic backgrounds, Related to Figures 2 and 3. **A)** Graph of actin polymerization over time. Two separate protein purifications of Arp2s-Arpc3B were compared. One protein batch was thawed after storage at -80°C, and the second was “fresh,” or never frozen. Rate of actin polymerization appeared the same between protein batches, suggesting protein quality is consistent between preps and not reduced due to freezing. **B)** Graph of maximum polymerization rates normalized to that of actin alone as in Fig. 2E. Different colors of the data points correspond to distinct batches of purified actin. Relates to panel A, showing independent protein purifications exhibit similar polymerization rates. Statistical test: an unpaired t-test with Welch’s correction. **C)** Image of AlphaFold 3 (Abramson et al., 2024) prediction of the *Drosophila* Arp2/3 complex with either Arp2s or Arp2L. The structure, predicted with actin monomers and ATP and Mg^2+^ as ligands, generated the expected active conformation. Predicted interaction between Arpc3A and the splice variants shown. Labeled residues are the interacting residues found in a previous *Bos taurus* Arp2/3 structure that form the binding interface (Ding et al., 2022). **D)** Crystal structure of CK-666 bound to bovine Arp2/3, PDB 3UKR (Baggett et al., 2012). The AlphaFold 3-predicted *Drosophila* Arp2L structure was aligned with the Arp2 fragment in the structure. The microexon is labeled in dark blue and is distal from the CK-666 position (magenta). The approximate position of CK-869 in Arp3 is indicated (red asterisk) based on the crystal structure PDB 3ULE (Hetrick et al., 2013). **E)** Left, crystal structure PDB 3ULE (Hetrick et al., 2013) with Arp3 shown bound to CK-869. Conservation between human and *Drosophila* Arp3 is mapped onto the structure. Conserved residues are in red and unconserved in blue, which are labeled with the human residue listed first. Right, crystal structure PDB 3UKR (Baggett et al., 2012) with Arp2 bound to CK-666. Conservation mapped as in left image. **F-H)** Graphs as in panel B. Complexes with Arp2s or Arp2L bound to either Arpc3A (“c3A”) or Arpc3B (“c3B”) were tested in polymerization assays with 100 µM CK-869, CK-666 or an equal volume of DMSO. Statistical test: a two-way ANOVA with a Tukey’s multiple comparisons post-hoc test comparing the means of each complex tested with or without drug. **I)** Plot showing the ratio of the maximum slope of a polymerization curve for Arp2L bound to Arpc3B plus 200 µM CK-666 or CK-869, divided by the maximum slope of the curve for Arp2L-Arpc3B plus DMSO. Statistical test: one-sample ratio t-test with the hypothetical value being 1. **J)** Left, schematic of crosses, in which flies homozygous for *Arp2L* in the *Arp2*-KO locus (“KO*-2L*”) or the second chromosome (“KO; *2L*”) were crossed. Right, superplot comparing the viability of progeny from the crosses depicted on the left. Percent of embryos that hatched within 48 hours were quantified. Each datapoint pertains to progeny laid in one day by the cross (average number of embryos quantified per datapoint: 174). Crosses with second chromosome transgenes (KO; *2s* and KO; *2L*) were done separately from the other crosses. Red and blue dots represent a separate experiment. Oregon-R flies (“WT”) were used as a control. Statistical test: one-way ANOVA with a Tukey’s multiple comparisons post-hoc test. KO-*2L* flies produced significantly less viable progeny compared to flies encoding *Arp2L* on the second chromosome, which was comparable to WT and *Arp2s*-expressing flies. **K)** Graph as in panel J. KO-*Arp2L* females were crossed to either WT (Oregon-R) or KO*-Arp2L* males, and KO-*Arp2L* males were crossed to WT females. KO-*Arp2L* females, not males, produced a reduced number of viable embryos. Crosses were allowed to lay embryos for four days, with each day’s progeny separately quantified for hatching within 48 hrs. Average number of embryos laid per day per cross: 125. **L)** Graph quantifying the number of eggs laid normalized per female. Oregon-R flies are “WT”. Statistical test as in J. Each datapoint represents a different cross with 10 or 9 females. Average number of embryos laid per cross: 185. **M)** Graph quantifying the percent of fertilized embryos in panel L produced by KO*-Arp2s* or KO*-Arp2L* females crossed to hemizygous KO*-Arp2s* or KO*-Arp2L* males. The number of embryos quantified per cross is indicated above each bar. Statistical test: chi-square test comparing fertilized to unfertilized raw counts. **N)** Graph quantifying the number of embryos in panel M that past cellularization in early embryogenesis. Statistical test as in panel M.

**Figure S3:**
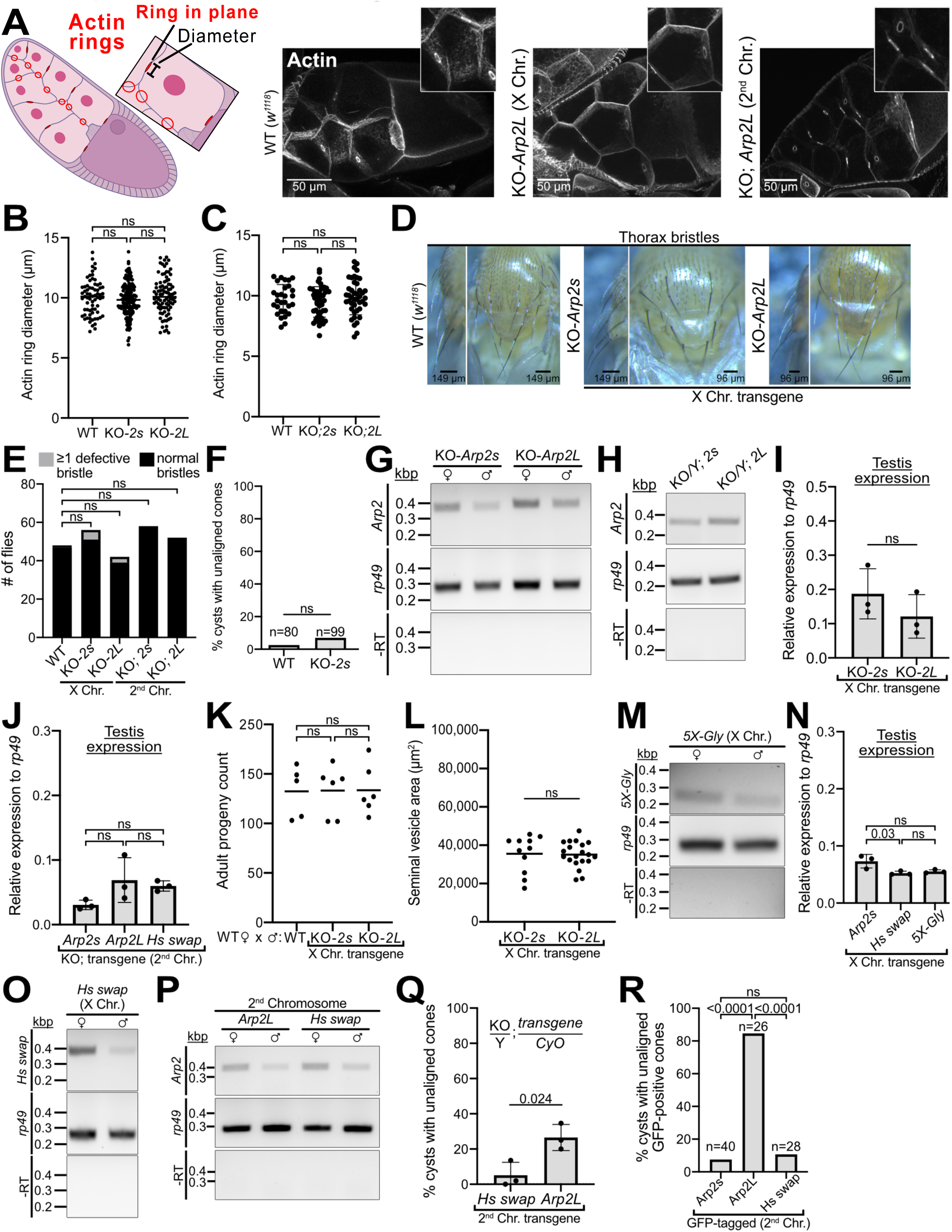
Tissue analyses of flies expressing single splice variants and characterization of the Arp2 chimeras, Related to Figures 3 and 4. **A)** Left, schematic of an egg chamber from an ovary. Actin rings in red are vestigial furrows critical in oogenesis (Hudson and Cooley, 2002). Right, representative images of actin-stained ovaries in WT (*w^1118^*) or *Arp2*-KO flies encoding *Arp2L* in the X or second chromosome. **B)** Graph quantifying the diameter of actin rings in stage 10 egg chambers, comparing ovaries from females encoding *Arp2s* or *Arp2L* in the *Arp2*-KO locus. WT flies: *w^1118^*. Statistical test: one-way ANOVA with a Tukey’s multiple comparisons post-hoc test. **C)** Graph as in panel B, comparing *Arp2*-KO females encoding *Arp2s* or *Arp2L* on the second chromosome. **D)** Representative images of bristles on the thorax (images on left are a side view to show curvature). Transgenic lines encode *Arp2s* or *Arp2L* in the X chromosome compared to *w^1118^*flies as WT. **E)** Quantification of the flies exhibiting one or more defective bristles (bent or wavy). Very few flies showed abnormal bristles. Statistical test: chi-square test comparing raw counts of normal versus abnormal bristles to that of *w^1118^*. **F)** Quantification of the percent of cysts with unaligned cones in WT (*w^1118^*) males and KO-*Arp2s* flies. The number of cysts analyzed (across ∼30 males per genotype) are noted above bars. Statistical test: Fisher’s exact test, showing no difference. KO-*Arp2s* values are from Fig. 3E **G)** RT-PCR analyses of *Arp2s* and *Arp2L* expression in whole females and whole males of KO-*Arp2s* or KO-*Arp2L* flies. *Rp49* served as a “housekeeping gene” to compare total cDNA quantity across the samples. The “-RT” (no reverse transcriptase) indicates no genomic DNA contamination. **H)** RT-PCRs as in panel G for *Arp2*-KO whole males encoding *Arp2s* or *Arp2L* on the second chromosome. Two copies of transgene present. **I-J)** Quantitative RT-PCR (RT-qPCR) analyses of *Arp2s* versus *Arp2L* expression in the testis of transgenic flies. Each datapoint corresponds to cDNA from 40 testes. Expression was calculated relative to *rp49*. Statistical tests: an unpaired t-test with Welch’s correction in panel I, and a one-way ANOVA with a Tukey’s multiple comparisons post-hoc test in panel J. **K)** Quantification of progeny from a male fertility test, in which KO-*Arp2s* or KO-*Arp2L* flies were crossed to WT females. Statistical test as in panel J. **L)** Quantification of the area of seminal vesicles from KO-*Arp2s* or *Arp2L* males. Seminal vesicles scale with number of sperm produced. Statistical test: unpaired t-test with Welch’s correction. No difference in area suggests similar sperm production. **M)** RT-PCRs of whole females or whole males showing expression of the *5X-Gly* chimera in the *Arp2*-KO background (see Fig. 4A), similar to panel G. **N)** RT-qPCR analyses of *Arp2s* versus the chimeras *Hs swap* and *5X-Gly* expression in the testis of transgenic flies. Each datapoint corresponds to cDNA from 40 testes. Expression was calculated relative to *rp49*. Statistical test: one-way ANOVA with a Tukey’s multiple comparisons post-hoc test. **O-P)** DNA gels showing RT-PCRs similar to panel G, verifying expression of the human swap (“Hs-swap”) chimera in whole females and whole males. Transgene in the *Arp2*-KO locus in panel O and the second chromosome in panel P. *Arp2L* in panel P shows comparable second-chromosome transgene expression. **Q)** Quantification of the percent of cysts with unaligned cones. Each datapoint represents a single cytological experiment quantifying total individualizing cysts across about 10 males per genotype. Statistical test: an unpaired t-test with Welch’s correction. **R)** Quantification of the percent of cysts with unaligned GFP-positive cones in males encoding GFP-tagged *Arp2s*, *Arp2L*, or the *human swap* chimera on the second chromosome in otherwise WT flies. Number of cysts analyzed are noted above bars. Statistical test: Fisher’s exact test with raw counts.

## Notes

### Competing Interest Statement

The authors have declared no competing interest.

### Summary of Updates

Following peer review, we have included additional in vitro and in vivo data that further support that the Arp2 microexon confers functional divergence of the Arp2/3 complex. In the revision, we find that the Arp2 microexon attenuates inhibition by Arp2/3 inhibitors, indicating intrinsic differences between Arp2 splice variants. Because of our new in vitro data, we have changed the title to "A microexon confers functional divergence of the actin nucleator Arp2/3" (previously, "A microexon in Arp2 alters tissue-specific Arp2/3-generated actin structures"). We also include additional analyses of fly tissues expressing only Arp2s versus Arp2L, as well as quantitative mRNA and protein expression analyses.

